# Essential Role of Placenta Derived Interferon Stimulated Gene 20 Against ZIKA Virus Infection

**DOI:** 10.1101/2021.01.13.426624

**Authors:** Jiahui Ding, Paulomi Aldo, Cai M. Roberts, Paul Stabach, Hong Liu, Yuan You, Xuemin Qiu, Jiwon Jeong, Anthony Maxwell, Brett Lindenbach, Demetrios Braddock, Aihua Liao, Gil Mor

## Abstract

Zika virus is a positive-sense single-stranded RNA virus, which can be transmitted across the placenta and leads to adverse effects on fetal development during pregnancy. The severity of these complications highlights the importance of prevention and treatment. However, no vaccines or drugs are currently available. In this study, we characterized the IFNβ-mediated antiviral response in trophoblast cells in order to identify critical components that are necessary for the successful control of viral replication; and determined whether we could use the components of the IFN-induced response as a replacement therapy for ZIKA viral infection during pregnancy. We identified and characterized interferon stimulated gene 20 (ISG20), playing a central role in controlling Zika infection in trophoblast cells, and successfully established a recombinant ISG20-Fc protein that effectively decrease viral titers in vitro and in vivo by maintaining its exonuclease activity and displaying immune modulatory functions. Therefore, rISG20-Fc is a promising anti-viral/immune modulatory approach for the control and prevention of RNA viral infections such as ZIKA virus.

## Introduction

During normal pregnancy, the placenta is capable of mounting a robust immune response and acting as a barrier to control different types of infections. Therefore, the placenta is the most important immune organ that protects the fetus from infectious pathogens, which pose major threats for fetal development (Mor et al., 2017). This protection is mediated by Pattern Recognition Receptors (PRR) that are expressed at the maternal-fetal interface, such as Toll Like Receptors (TLRs) and the expression of anti-viral factors, such as type I Interferon attributes a central role in viral infection clearance (Cardenas et al., 2011, Koga and Mor, 2010). However, in some cases, like during ZIKA virus (ZIKV) infection, this protective barrier is breached and leads to serious pathologic consequences for the fetus development.

ZIKV is a positive-sense single-stranded RNA virus, which belongs to the *Flavivirus* genus in the *Flaviviridae* family (Dick et al., 1952). Infection with ZIKV during pregnancy leads to adverse pregnancy outcomes, including preterm birth, stillbirth, and congenital Zika syndrome (CZS) (Sadovsky et al., 2016, Schwartz, 2017b, Shan et al., 2016). Infants with CZS may show microcephaly, abnormal brain development, limb contractures, eye abnormalities and other neurologic manifestations (Rasmussen et al., 2016, Karimi et al., 2016, Johansson et al., 2016). Moreover, a recent study shows that infants exposed to ZIKV infection in utero showed neurodevelopmental delays as toddlers, despite having “normal” brain imaging and head circumference at birth (Honein et al., 2020). This highlights the long-term effects of ZIKV infection on offspring development and further implicates the need to protect mother and fetus against ZIKV infection during the early stage of pregnancy.

As of July 2019, ZIKV has been documented in 87 countries and territories’ (Brady et al., 2019, Franca et al., 2016, Organization, 2018, Organization, 2019). In the Americas, the ZIKV outbreak peaked in 2016, but the incidence subsequently declined during 2017 and 2018. However, in 2018, there were still a total of 31,587 suspected, probable, and confirmed cases of ZIKV disease reported (Organization, 2019, Pattnaik et al., 2020, Honein et al., 2020). All areas with prior ZIKV infection have the potential for re-emergence and re-introduction. From the facts above, it is clear that ZIKV infection poses a high risk for human reproduction health and results in a massive economic burden and workload on the healthcare systems, which calls for the rapid development of safe and efficacious vaccines and therapeutics.

Although major efforts have been invested on the development of vaccines and anti-ZIKA drugs(Pattnaik et al., 2020), no one has been materialized and the number of ZIKV-infected patients continues to grow; therefore there is a need for a better understanding on the cellular mechanism that provide effective protection during infection that could lead to the development of more effective therapeutic approaches that can prevent ZIKV-induced developmental abnormalities in fetus.

Type I Interferon (IFN) signaling plays a crucial role in controlling ZIKV replication and pathogenesis (Aliota et al., 2016, Lazear et al., 2016, Yockey et al., 2016). As demonstrated in mice models lacking type I IFN signaling, these IFNAR1^−/−^ mice were more susceptible to ZIKV infection and developed neurological disease (Lazear et al., 2016, Miner et al., 2016a, Racicot et al., 2017). Furthermore, in our previous study, we demonstrated the specific fetal/placental derived type I IFN signaling plays a critical role in the prevention of fetal viral infection, and is the key for ensuring maternal survival (Racicot et al., 2017, Kwon et al., 2018, Aldo et al., 2016b). This protective effect depends on the integrity of the IFNβ/IFNAR1 pathway and their downstream Interferon stimulated genes (ISGs).

Type I IFNs, including IFN-α, IFN-β, IFN-∊, IFN-κ, and IFN-ω, are the primary IFNs that are generated in most cell types during infections. The canonical IFN response will be initiated upon recognition of viral genomes/transcripts by RNA-sensing helicases, such as the retinoic acid-inducible gene I (RIG-I) and melanoma-associated differentiation antigen 5 (MDA5) (Kell and Gale, 2015, Said et al., 2018). After binding viral RNA, RIG-1/MDA5 recruits the adaptor mitochondrial antiviral-signaling protein (MAVS) to trigger the phosphorylation of the antiviral transcription factors IFN regulatory factor 3 (IRF3) and NF-κB, which leads to the induction of type I IFN production (Pichlmair and Reis e Sousa, 2007, Zevini et al., 2017). Secreted type I IFN binds to the heterodimeric transmembrane receptor IFNAR, consisting of IFNAR1 and IFANR2 chains, and signals through the JAK/STAT pathway to induce the expression of hundreds of ISGs (Levy et al., 2011, Platanias, 2005). These ISGs exert a potent anti-viral response on specific steps of the viral life cycle and inhibit virus replication and shedding (Schneider et al., 2014). An early step in the cellular anti-viral response is to target the viral RNA before it is able to replicate and produce new viral particles to infect neighboring cells. This process is accomplished through the expression of specific exonucleases (Schneider et al., 2014).

The objectives of this study were two-fold: 1) to characterize the IFNβ-mediated antiviral response in trophoblast cells in order to identify critical components that are necessary for the successful control of viral replication; and 2) to determine whether we could use the components of the IFN-induced response as a replacement therapy for ZIKV infection during pregnancy in cases where its function has being affected. We found that ISG20, an interferon-inducible 3’-5’ exonucleases with potent anti-viral activity against different viruses (Espert et al., 2005, Jiang et al., 2008, Liu et al., 2017, Qu et al., 2016, Zhang et al., 2007) is one of the earlier ISGs induced by a protective IFNβ-mediated response. Deletion of ISG20 in trophoblast cells decreases their capacity to control ZIKV replication and render these cells susceptible to the infection.

In this study, we report the characterization of ISG20 expression and function during ZIKV infection in trophoblast cells and its central immune modulatory function. Use of a recombinant ISG20 protein, which conserves its RNase activity, restores protection against ZIKV by inhibiting ZIKV replication in vitro and in vivo. Our findings highlight the role of ISG20 as one of the key ISGs responsible for inhibiting ZIKV infection and demonstrate its potential application for the treatment/prevention of ZIKV infections during pregnancy.

## Results

### 1. ZIKV infection induces IFNβ and ISGs mRNA expression in first-trimester trophoblast cells

Our first objective was to characterize the cellular components involved during an effective anti-viral response elicited by trophoblast cells exposed to ZIKV. We used a human first-trimester trophoblast cell line (Sw.71), which has been well characterized in previous studies (Aldo et al., 2007, Straszewski-Chavez et al., 2009, Aldo et al., 2010), and primary cultures of human first-trimester trophoblast cells.

Both cell types were infected with Cambodia ZIKV (MOI=2) for 1 h, refreshed with regular media and monitored for 48 h to determine changes in cell growth and survival. During the infection, Sw.71 cells and primary cultures of first-trimester trophoblast infected with ZIKV did not show any apoptosis-related morphological changes or disturbance on cell growth when compared to the control, which suggests that trophoblast cells are capable of controling ZIKV infection. This observation was confirmed when we evaluated viral titers at different time points post infection by qRT-PCR. We detected ZIKV titers in Sw.71 at 24 h post infection (h.p.i.), with higher levels at 48 h.p.i. (Fig.1A). Sw.71 cells were able to control the infection as shown by significant decrease on viral titers at 72 h.p.i. (Fig.1A). In human primary trophoblast cell cultures (HPC), we observed that ZIKV titers peaked at 72h.p.i but started to decrease after 96 h.p.i. (Fig.1A). Due to the technical limitations of maintaining primary cultures for longer time, we were not able to further evaluate the decrease on viral titers at later times (more than 96 h.p.i.), but as indicated above we did not observe any sign of cell death in the primary trophoblast cell cultures.

**Figure 1.**
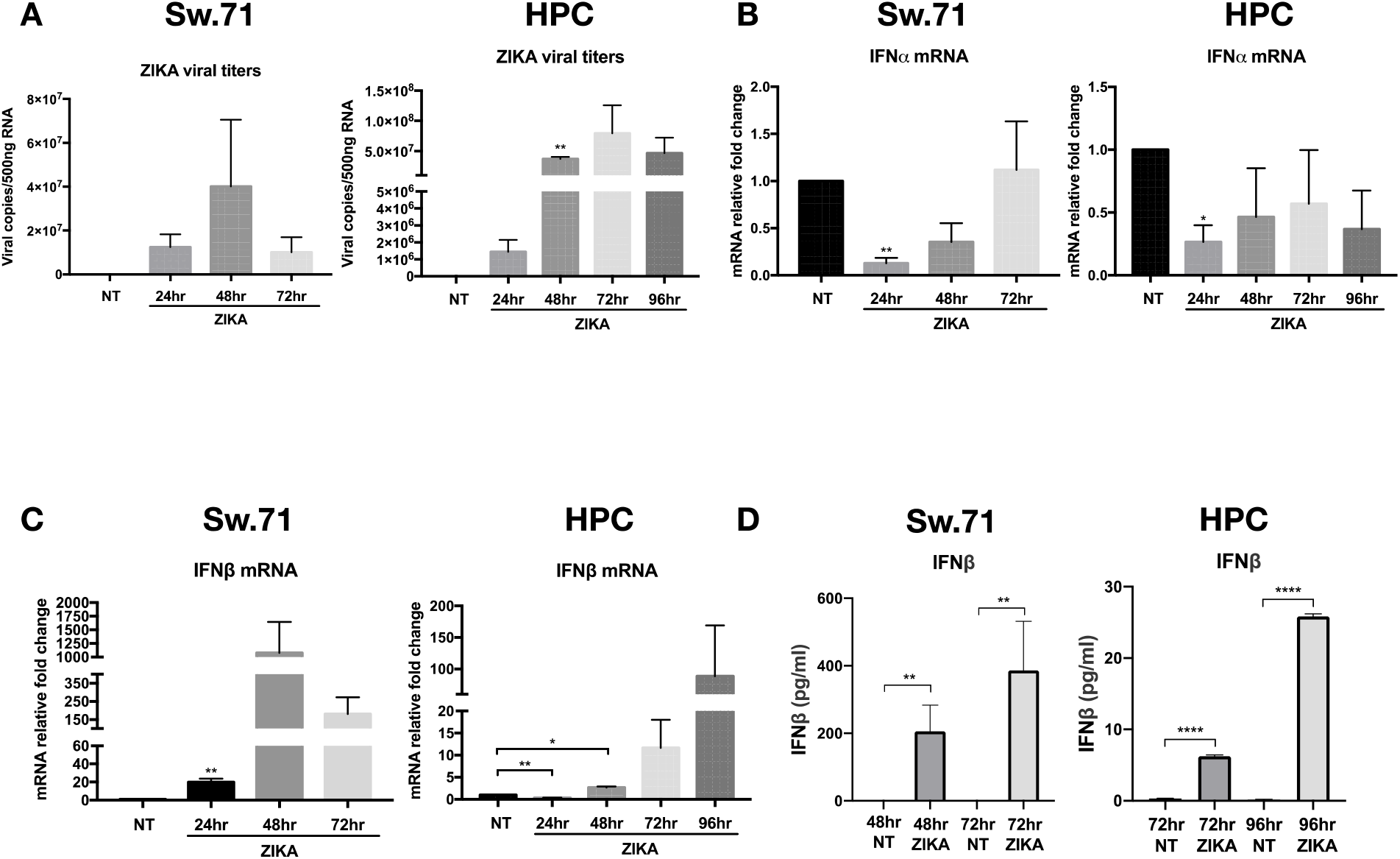
ZIKV infection induces type I IFNβ expression in human first-trimester trophoblast cell line Sw.71 and human primary cultures. Human first-trimester trophoblast cell line Sw.71 and human primary culture trophoblast cells were infected with ZIKV (MOI=2) for 1 h and refreshed with regular media over time. RNA was collected for measuring viral titers and gene expression by qRT-PCR. Data represent as mean ± SEM from three independent experiments. \**p* < 0.05, \*\**p* < 0.01, \*\*\*\**p* < 0.0001. NT, no treatment group; HPC, human primary culture. **(A)** ZIKV titers in Sw.71 and HPC trophoblasts. Note that ZIKV titers in Sw.71 increased in the first 24 hours post infection (h.p.i.) and reached the peak at 48 h.p.i., then decreased at 72 h.p.i. While in HPC, ZIKV titers went up until 72 h.p.i., and decreased at 96 h.p.i. **(B)** IFNα is not activated in response to ZIKV infection in trophoblast cells. IFNα mRNA expression was inhibited in the first 24 h.p.i. and maintained at the basal level during ZIKA infection. **(C)** ZIKV infection induced a time-dependent increase in IFNβ mRNA expression in Sw.71 and HPC. Note that at 24 h.p.i, IFNβ expression was inhibited in HPC and then increased afterwards. **(D)** Expression levels of secreted IFNβ detected in trophoblast supernatants by ELLA assay. Supernatants were collected from ZIKV-infected and control trophoblast cultures at different times, and IFNβ protein secretion was quantified by ELLA assay. Note the increase of secreted IFNβ in the ZIKV-infected supernatant in a time-dependent manner.

To understand the anti-viral response generated by trophoblast cells to ZIKV infection, the type I interferon response was first characterized in Sw.71 and HPC infected with ZIKV (MOI=2) over different time points post ZIKV infection. Interestingly, we observed the inhibition of IFNα mRNA expression by ZIKV infection (Fig.1B), but a time-dependent increase of IFNβ mRNA (Fig.1C), which was consistent with the time-dependent pattern of ZIKV titers. IFNβ mRNA expression in Sw.71 increased in the first 24 h.p.i., peaked at 48 h.p.i., and further decreased 72 h.p.i. (Fig.1C). In HPC the IFNβ response followed similar pattern as the viral infection; IFNβ mRNA increased at 48 h.p.i. and remained high at 72 h.p.i. and 96 h.p.i. (Fig.1C). Although the response times were different between the cell line and the primary cultures, both cell types showed significant increase of IFNβ expression in response to ZIKV infection (Fig. 1).

To validate the observed increase of IFNβ mRNA expression and its potential function following ZIKV infection, we tested the presence of secreted IFNβ protein in the supernatant of Sw.71 and HPC trophoblast cells at different time points post ZIKV infection. We did not detect secreted IFNβ in the supernatant of control non-infected Sw.71 cells, however, a substantial concentration of secreted IFNβ protein was detected in the supernatant of ZIKV-infected Sw.71 cells at 48 and 72 h.p.i. (Fig.1D). While in HPC, IFNβ protein was undetectable in the supernatant both from non-infected and ZIKV-infected cells at 24 and 48 h.p.i. (data not shown), a significant increase of secreted IFNβ protein was observed at 72 and 96 h.p.i (Fig.1D). Therefore, these findings confirm that trophoblast cells can recognize ZIKV infection and initiate a type I IFNβ response which is efficient to control viral infection.

### 2. IFNβ promotes ISGs expression in trophoblast cells in response to ZIKV infection

Following its secretion from cells, IFNβ mediates its anti-viral activity by inducing the expression of Interferon stimulated genes (ISGs) (Schneider et al., 2014). Thus, we evaluated the mRNA expression of the anti-viral ISGs: *ISG20, MX1, OAS1, ISG15, CH25H, TRIM22, Tetherin*, and *Viperin* in Sw. 71 and HPC trophoblast cells following ZIKV infection. Similar to IFNβ expression, we observed a significant increase of the mRNA expression for all these ISGs at 48 h.p.i. followed by decreased expression levels at 72 h.p.i. in Sw.71 cells infected with ZIKV (Sup. Fig.1). HPC showed similar response although at different times, with a peak of ISGs mRNA expression at 96 h.p.i.; which correlated with the time of IFNβ peak expression and the observed decrease on viral titers (Sup. Fig.2).

To determine whether these ZIKA-induced ISGs are downstream of IFNβ, Sw.71 cells were treated with increasing doses of IFNβ (3, 30, 300 ng/ml) for 8 h, and ISGs mRNA expression was determined by qRT-PCR. Our data shows that IFNβ was able to induce the expression of these ISGs in a dose-dependent manner (Sup. Fig.3). These results suggest that IFNβ-induced ISGs expression in trophoblast cells represent the first line of response to ZIKV infection. Interestingly, we did not observe a similar IFNβ/ISGs response in trophoblast cells infected with Herpes Simplex Virus-2 (HSV-2) (Data not shown), suggestive of the specific nature of IFNβ response to ZIKV infection.

### 3. Induction of ISG20 expression by IFNβ in response to ZIKV infection

An early step in the cellular anti-viral response is to target the viral RNA before it is able to replicate and produce new viral particles. This process is accomplished through the expression of exonucleases that specifically degrade viral RNA (Schneider et al., 2014). In our screening of the anti-viral ISGs, we observed the induction of ISG20 in human first-trimester trophoblast cells infected with ZIKV (Sup.Fig.1 and 2). ISG20 functions as an interferon-inducible 3’-5’ exonuclease and has been shown to exert a potent antiviral activity against different viruses (Espert et al., 2005, Jiang et al., 2008, Liu et al., 2017, Qu et al., 2016, Zhang et al., 2007). Therefore, we pursued the characterization of ISG20 because of its early potential interruptive function with the viral cycle.

First, we confirmed that ISG20 expression is directly regulated by IFNβ. Thus, Sw.71 or HPC were treated with increasing concentrations of IFNβ (3, 30, 300 ng/ml) for 8 h and ISG20 mRNA expression was quantified by qRT-PCR. Our data showed that IFNβ was a very strong inducer of ISG20 mRNA expression in trophoblast cells (Sw.71 and HPC) and its effect was dose-dependent (Fig.2A). Furthermore, the induction of ISG20 by IFNβ was early (Fig.2B), where we observed a significant increase of ISG20 mRNA expression as early as 2 h following exposure to IFNβ (300 ng/ml) and was further enhanced after 16 h and 24 h (Fig.2B).

**Figure 2.**
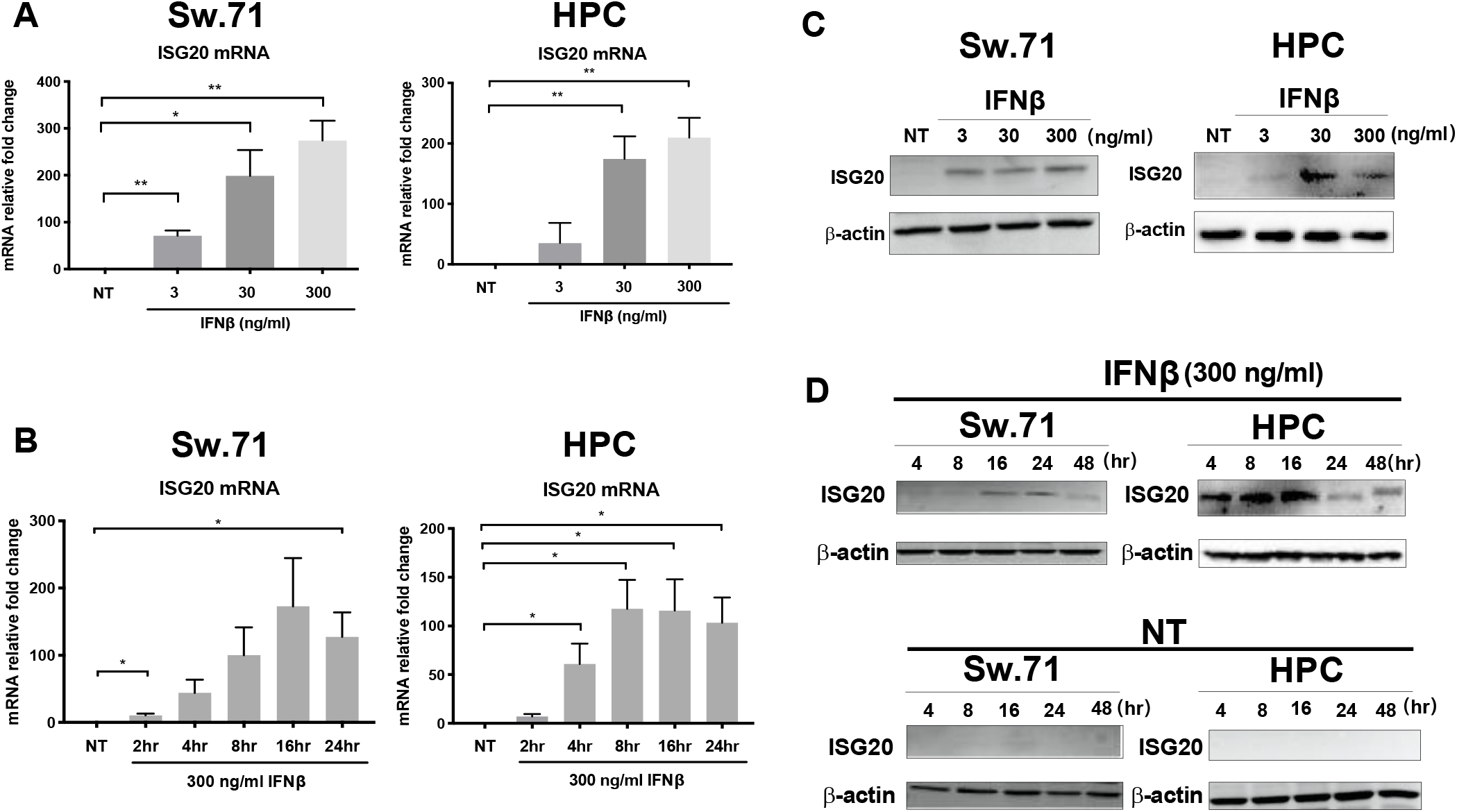
IFNβ promotes ISG20 mRNA and protein expression in a dose and time-dependent manner in trophoblast cells. **(A)** Sw.71 and HPC cells were treated with different doses of IFNβ (3, 30, 300 ng/ml) for 8 h and RNA were collected for determining ISG20 mRNA expressions by qRT-PCR. Note the increase of ISG20 mRNA expression in a dose-dependent manner. **(B)** Sw.71 and HPC cells were treated with 300ng/ml IFNβ over time and RNA was collected for determining ISG20 mRNA expressions by qRT-PCR. Note the increase of ISG20 mRNA expression in a time-dependent manner. **(C)** Sw.71 and HPC cells were treated with different doses of IFNβ (3, 30, 300 ng/ml) for 24 h and protein was collected for determining ISG20 protein expressions by western blot. Note the increase of ISG20 protein expression in a dose-dependent manner. **(D)** Sw.71 and HPC cells were treated with 300ng/ml IFNβ over time and protein was collected for determining ISG20 protein expressions by western blot. Note that there was no ISG20 protein expression in the no treatment group in both Sw.71 and HPC, only after IFNβ treatment, ISG20 protein expression exhibited a time-dependent manner. Data represent as mean ± SEM from three independent experiments. β-actin served as a loading control for western blot. \**p* < 0.05, \*\**p* < 0.01. NT, no treatment group; HPC, human primary culture.

Next, we determined if the changes in ISG20 mRNA were translated into protein expression. Trophoblast cells were treated with increasing concentrations of IFNβ for 24 h and protein expression was determined by western blot analysis. Interestingly, we found no protein expression in non-stimulated trophoblast cells (Fig.2C and D); even though they have high basal mRNA levels as shown by the Cq value ranging from 23 to 26 (data not shown). However, IFNβ was able to induce ISG20 protein expression in a dose and time-dependent manner in both Sw.71 and HPC (Fig.2C and D). Intriguingly, this protein expression is tightly regulated. We saw ISG20 protein expression as early as 4 h post IFNβ treatment and increased up to 16 h in HPC. Afterwards, ISG20 protein expression decreased to almost basal levels (Fig.2C and D). This data demonstrates that ISG20 transcription and translation are both inducible by IFNβ in first-trimester trophoblast cells.

### 4. Poly(I:C) induces IFNβ and ISG20 expression in trophoblast cells

Our next objective was to investigate ISG20 expression in response to a general viral RNA. Therefore, we first used Polyinosinic: polycytidylic acid (Poly I:C), which is a synthetic double-stranded RNA that can mimic viral infection when applied in vitro and in vivo (Fortier et al., 2004). We tested the hypothesis that ISG20 expression could be induced by Poly(I:C) treatment in trophoblast cells. We treated Sw.71 and primary trophoblast cells with different doses of Poly(I:C) (0.25, 2.5, 25 μg/ml). IFNβ and ISG20 mRNA and protein expression were determined by qRT-PCR and western blot, respectively. Poly(I:C) treatment induced IFNβ and ISG20 mRNA expression in a dose-dependent manner in Sw.71 and HPC trophoblast cells (Fig.3A). At the protein level, Poly(I:C) also induced ISG20 protein expression in a dose-dependent manner in both cell types (Fig.3B). HPC trophoblasts seemed to be more sensitive to Poly(I:C) treatment since we were able to detect increased levels of ISG20 protein expression when cells were treated with Poly(I:C) at its lowest concentration (0.25 μg/ml) (Fig.3B).

**Figure 3.**
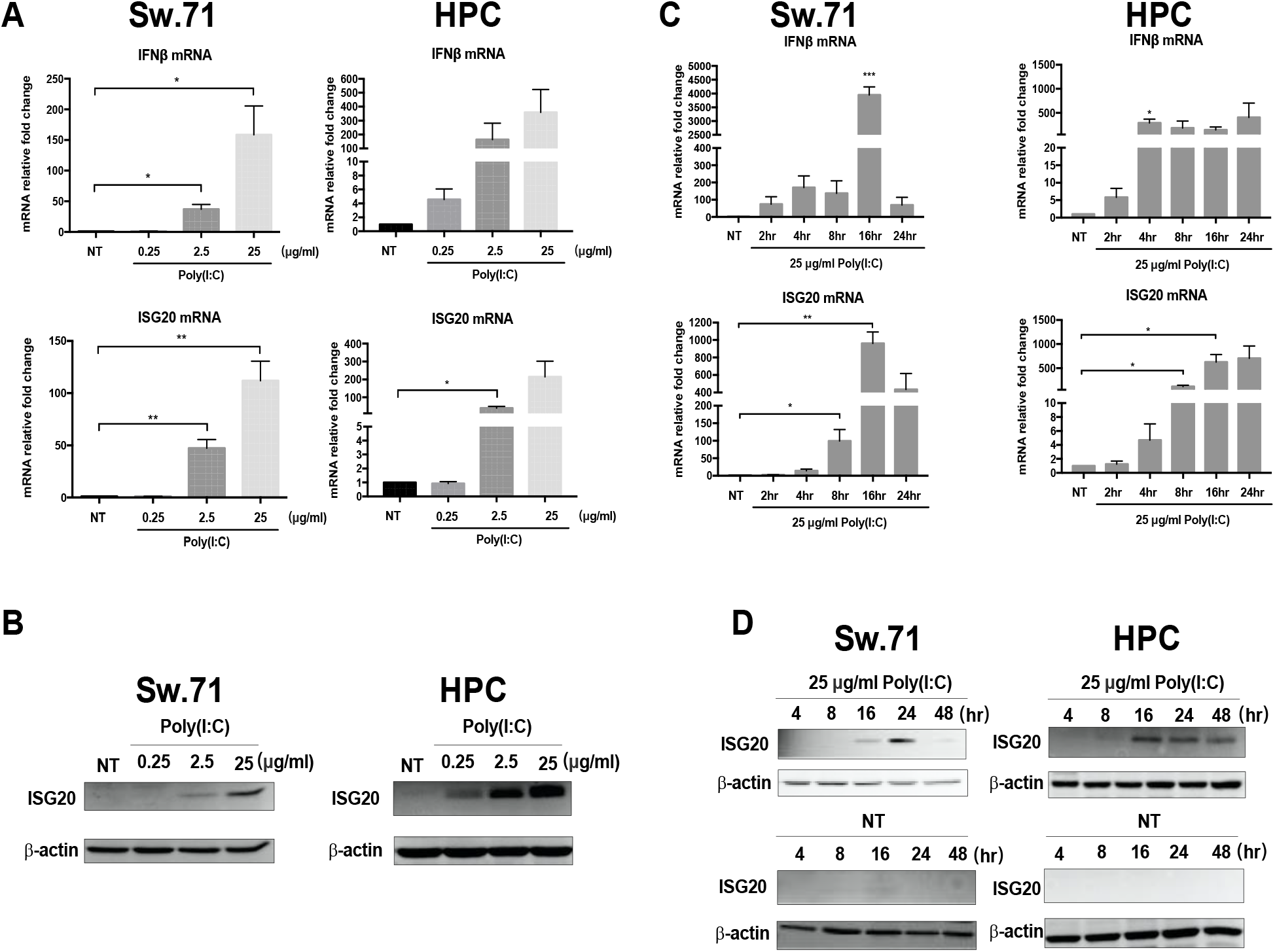
Poly(I:C) treatment leads to a dose and time-dependent increase of IFNβ and ISG20 expression in trophoblast cells. **(A)** Sw.71 and HPC cells were treated with different doses of Poly(I:C) (0.25, 2.5, 25 μg/ml) for 8 h and RNA was collected for determining ISG20 mRNA expressions by qRT-PCR. Note the increase of IFNβ and ISG20 mRNA expression by Poly(I:C) in a dose-dependent manner. **(B)** Sw.71 and HPC cells were treated with different doses of Poly(I:C) (0.25, 2.5, 25 μg/ml) for 24 h and protein was collected for determining ISG20 protein expressions by western blot. Note the increase of ISG20 protein expression in a dose-dependent manner. **(C)** Sw.71 and HPC cells were treated with 25 μg/ml Poly(I:C) over time and RNA was collected for determining IFNβ and ISG20 mRNA expressions by qRT-PCR. Note the increase of IFNβ and ISG20 mRNA expression in a time-dependent manner. **(D)** Sw.71 and HPC cells were treated with 25 μg/ml Poly(I:C) over time and protein was collected for determining ISG20 protein expressions by western blot. Note again that there was no ISG20 protein expression in the no treatment group in both Sw.71 and HPC, only after Poly(I:C) treatment, ISG20 protein expressed in a time-dependent manner. Data represent as mean ± SEM from three independent experiments. β-actin served as a loading control for western blot. \**p* < 0.05, \*\**p* < 0.01, \*\*\**p* < 0.001. NT, no treatment group; HPC, human primary culture.

Based on these findings, we then determined the earlier time when Poly(I:C) can promote IFNβ and ISG20 expression in trophoblast cells. Accordingly, Sw.71 and primary trophoblast cells were treated with Poly(I:C) (25 μg/ml) and cell pellets were collected at 2, 4, 8, 16, 24 h post treatment for mRNA evaluation and 4, 8, 16, 24, 48 h post treatment for protein evaluation. As we can see in figure 3C, Poly(I:C) treatment in Sw.71 cells induced IFNβ mRNA expression as early as 2 h post treatment reaching higher levels at 16 h and decreasing at 24 h (Fig.3C). ISG20 mRNA expression followed similar pattern of expression as IFNβ, although we detected the earliest increase at 4 h post treatment, and a major increase at 16 and 24 h (Fig.3C). In HPC, we observed a similar early response to Poly(I:C) for IFNβ and ISG20. However, contrary to Sw.71 cells, the mRNA levels remained higher even at 24 h post treatment (Fig.3C). We saw a comparable response in protein expression when HPC were treated with Poly(I:C). As indicated above, trophoblast cells do not express ISG20 protein in basal conditions. However, following treatment with Poly(I:C) (25 μg/ml), we were able to detect ISG20 protein expression as early as 16 h post treatment (Fig.3D). Altogether, this data suggests that viral RNA and Poly(I:C) can be sensed by trophoblasts cells, which will lead to the induction of a Type I IFNβ response and expression of ISG20.

### 5. Role of ISG20 in controlling ZIKV infection in trophoblast cells

Next, we sought to evaluate if trophoblasts would elicit a similar ISG20 response to ZIKV infection as the one observed with Poly (I:C). Firstly, we infected Sw.71 with ZIKA virus (MOI=2) for 1 h and collected cell pellets at 24, 48, 56 and 72 h.p.i., and determined ISG20 protein expression by western blot. Our results confirmed that ZIKV infection induced ISG20 protein expression (Fig. 4A) in a time-dependent manner, which was correlated to the increase in ISG20 mRNA (Sup. Fig.1). Moreover, ISG20 protein expression was detected at 24, 48 and 56 h.p.i. and decreased at 72 h.p.i. (Fig.4A).

**Figure 4.**
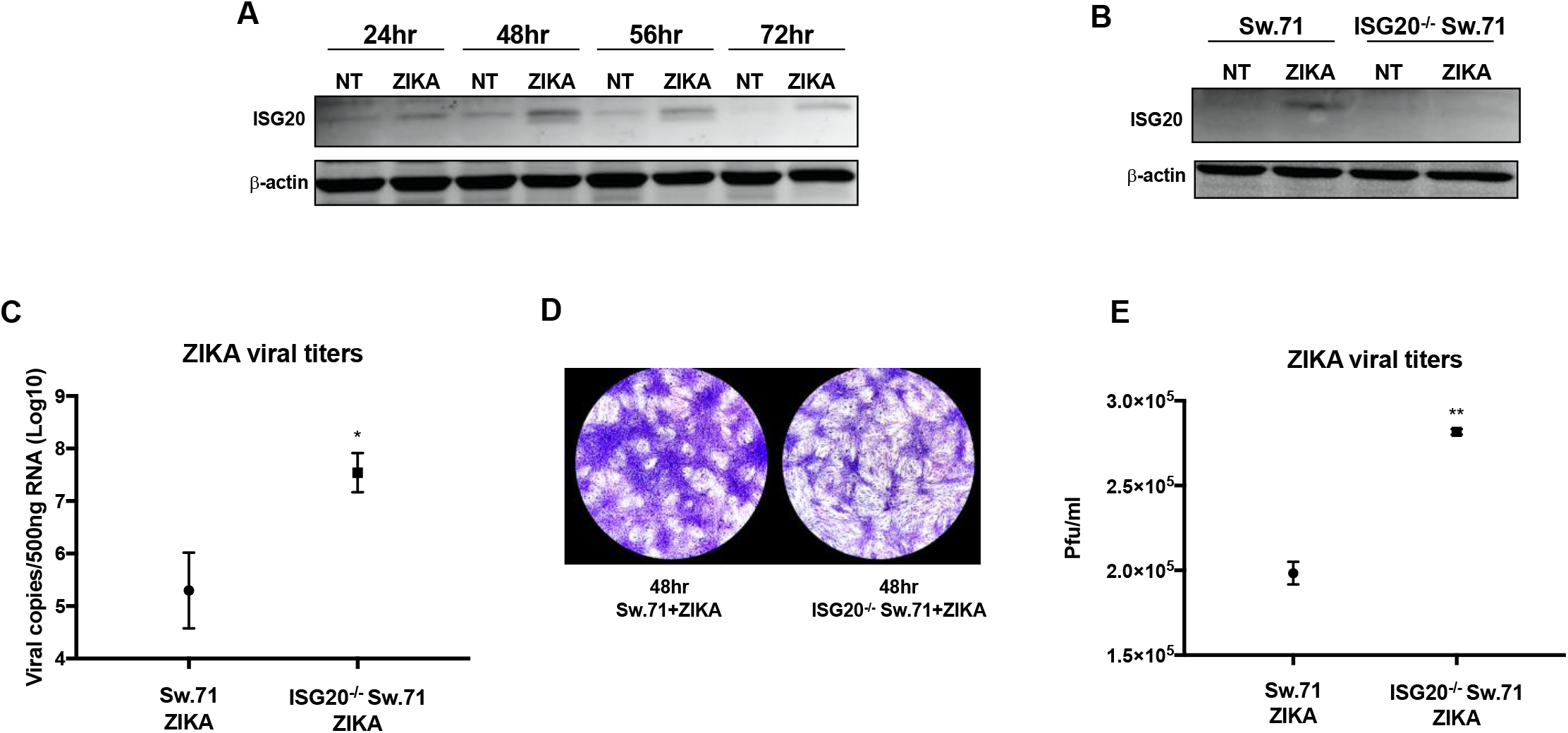
Lack of ISG20 in trophoblast cells leads to more ZIKV replication and viral shedding. **(A)** ISG20 protein expression was increased in ZIKV infection. Sw.71 cells were infected with ZIKV (MOI=2) over time, and proteins were collected for western blot analysis. **(B)** No ISG20 protein expression in ISG20^−/−^ Sw.71 in ZIKV infection. Sw.71 and ISG20^−/−^ Sw.71 cells were infected with ZIKV (MOI=2) for 1 h and refreshed with regular media for 48 h, and proteins were collected for western blot analysis. **(C)** Higher ZIKA titer was shown in ISG20^−/−^ Sw.71 cells. Sw.71 and ISG20^−/−^ Sw.71 cells were infected with ZIKV (MOI=2) for 1 h and refreshed with regular media for 48 h, and RNA was collected for determining the viral titer and gene expression by qRT-PCR. **(D)** More viral shedding in ZIKV-infected ISG20^−/−^ Sw.71 cell culture supernatant. Supernatant from ZIKV-infected Sw.71 and ISG20^−/−^ Sw.71 cells were collected and plaque assay was performed using Vero cells. A representative plaque assay picture is presented. Note that more plaques were formed in Vero cells by incubating with ZIKV-infected ISG20^−/−^ Sw.71 cell culture supernatant. **(E)** Plaques were counted and averaged for the quantification of the viral titers. Data represent as mean ± SEM from three independent experiments. β-actin served as a loading control. \**p* < 0.05, \*\**p* < 0.01.

To further elucidate the specific role of ISG20 during ZIKV infection, we established a trophoblast cell line lacking ISG20 (ISG20^−/−^ Sw.71) using the CRISPR-Cas9 system. The validation of ISG20^−/−^ Sw.71 was evidenced by the lack of ISG20 protein expression following Poly(I:C) treatment (Sup Fig.4). We then infected wild type (wt) Sw.71 and ISG20^−/−^ Sw.71 with ZIKA virus (MOI=2) for 1 h and incubated with refreshed growth media for 48 h. Afterwards, RNA and protein were collected to determine viral titers by qRT-PCR and protein expression by western blot. In contrast to wt Sw.71 cells, which showed ISG20 protein expression following ZIKV infection, no detectable ISG20 protein expression was observed in ISG20^−/−^ Sw.71 (Fig.4B). More importantly, ISG20^−/−^ Sw.71 exhibited significant higher viral titer levels when compared to wt Sw.71 (Fig.4C); suggesting that the lack of ISG20 rendered trophoblast cells more susceptible to ZIKV infection.

Subsequently, we evaluated whether the lack of ISG20 could have an impact in the process of ZIKV shedding. Vero cells were cultured in the presence of supernatants collected from 48h ZIKV-infected wt Sw.71 and ISG20^−/−^ Sw.71 cells, and ZIKV shedding was determined by plaque assay. As shown in Fig. 4D and E, more plaques were formed in Vero cells exposed to supernatants from ZIKV-infected ISG20^−/−^ Sw.71 group compared to supernatants from the ZIKV-infected wt Sw.71 group.

To further understand the mechanism responsible for the increased viral titer in ISG20^−/−^ Sw.71, we assessed whether the lack of ISG20 would affect the expression of other ISGs necessary for the anti-ZIKV response. First, we assessed the levels of IFNβ expression after ZIKV infection and observed that, although the levels of induction were different, wt Sw.71 cells and ISG20^−/−^ Sw.71 cells maintained the capacity to recognize and produce IFNβ (Fig.5A). Next, we evaluated the mRNA expression of several other anti-viral ISGs (*MX1, OAS1, ISG15, CH25H, TRIM22, Tetherin* and *Viperin)* in ZIKV-infected wt Sw.71 and ISG20^−/−^ Sw.71 cells. As shown in figure 5B, both cell lines showed an increased expression of anti-viral ISGs, while the only major difference between the two cell lines was the lack of ISG20 in the ISG20^−/−^ Sw.71 trophoblast cells (Fig.5C). Interestingly, although we observed increased mRNA levels for the tested ISGs in ISG20^−/−^ Sw.71 trophoblast cells, the fold changes were not as robust as those observed in wt Sw.71 (Fig.5B), which suggests that ISG20 may play a role in regulating the expression of other ISGs during ZIKV infection.

**Figure 5.**
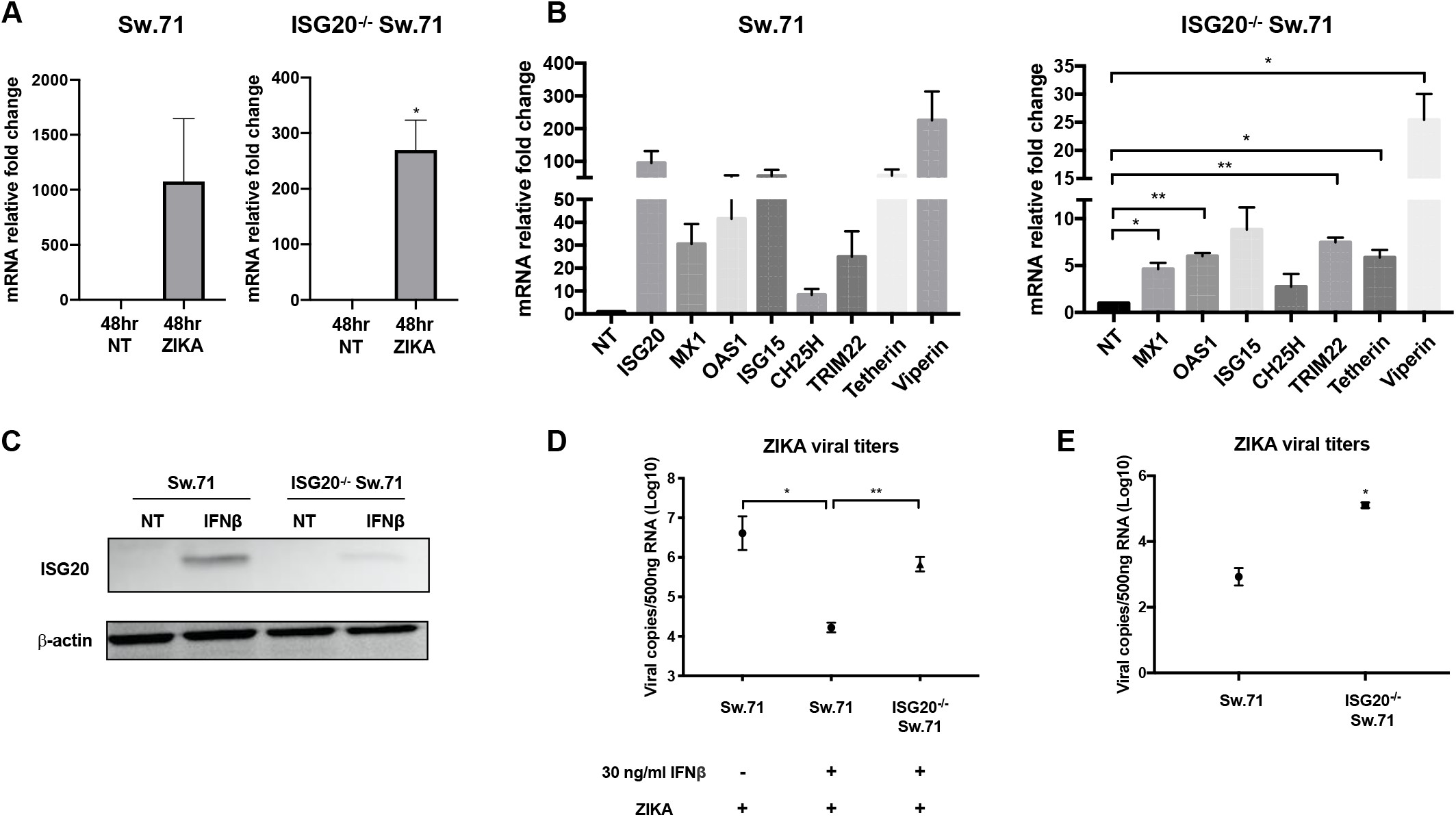
Lack of ISG20 in trophoblast cells weakens the protection provided by IFNβ against ZIKV infection. **(A)** Induction of IFNβ mRNA in response to ZIKV infection in Sw.71 and ISG20^−/−^ Sw.71 trophoblast cells. Sw.71 and ISG20^−/−^ Sw.71 cells were infected with ZIKV (MOI=2) for 1 h and refreshed with regular media for 48 h, and RNA was collected for qRT-PCR. Note that ISG20^−/−^ Sw.71 cells maintain the capacity to recognize and produce IFNβ. **(B)** ISGs mRNA expression stimulated by IFNβ in Sw.71 and ISG20^−/−^ Sw.71 trophoblast cells. Sw.71 and ISG20^−/−^ Sw.71 cells were treated with 30 ng/ml IFNβ for 24 h, and RNA was collected for determining the ISGs gene expression by qRT-PCR. Note that less fold changes were shown in ISG20^−/−^ Sw.71 cells after IFNβ treatment. **(C)** No ISG20 protein expression in ISG20^−/−^ Sw.71 cells after IFNβ treatment. Sw.71 and ISG20^−/−^ Sw.71 cells were treated with 30 ng/ml IFNβ for 24 h, and proteins were collected for western blot analysis. **(D)** IFNβ pre-treatment significantly prevented trophoblast cells from ZIKV infection, however, this protection was evidently attenuated due to lack of ISG20. Sw.71 and ISG20^−/−^ Sw.71 cells were pre-treated with or without 30 ng/ml IFNβ for 24 h, followed by ZIKV infection (MOI=2) for 1 h and refreshed with regular media for 24 h, and RNA was collected to determine the viral titers by qRT-PCR. **(E)** More viral shedding in ZIKV-infected ISG20^−/−^ Sw.71 cell culture with IFNβ pre-treatment. Supernatant from ZIKV-infected Sw.71 and ISG20^−/−^ Sw.71 cells with IFNβ pretreatment were collected and infected HESC for 1 h and refreshed with growth media for 24 h, RNA was collected to determine the viral titers by qRT-PCR. Data represent as mean ± SEM from three independent experiments. β-actin served as a loading control for western blot. \**p* < 0.05, \*\**p* < 0.01.

We then evaluated the response to ZIKV infection in IFNβ pre-treated trophoblast cells, and showed that treatment of wt Sw.71 with IFNβ (30 ng/ml) was able to increase ISG20 protein expression and significantly decreased ZIKV titers (Fig.5C and D). However, similar pre-treatment with IFNβ in ISG20^−/−^ Sw.71 failed to induce ISG20 and had minimal effect on controlling ZIKV titers (Fig.5C and D). Similarly, IFNβ pre-treatment of ISG20^−/−^ Sw.71 did not decrease viral shedding, which is shown by the significantly higher ZIKV titer of HESC cells that were exposed to supernatants from ISG20^−/−^ Sw.71 (Fig.5E). In summary, this data demonstrates that ISG20 is an early and critical component of the anti-ZIKV response, which functions by inhibiting ZIKV replication and dissemination in trophoblast cells.

### 6. Characterization of the anti-viral effect of recombinant ISG20

We hypothesized that ISG20 could be used as an anti-viral treatment by blocking early stages of viral replication. To test this hypothesis, we developed a recombinant form of ISG20 by cloning the full-length of human ISG20, a linker (Gly-Ser-Gly-Ser-Gly), and the human Fc domain into pcDNA4 vector. We also added a cell secretion signal and a His tag for purification. The protein structure is shown in Figure 6A and includes the following regions: 1) ISG20 sequence (Pink); 2) Linker (Orange); 3) Fc domain (White); 4) Tev sequence, cleavage between Q and G (Cyan); 5) secretory signal sequence (ENPP7) (Red); 6) His Tag (Green). We transfected this plasmid into Chinese hamster ovary (CHO) cells and selected for plasmid integration by growing the cells in the presence of Zeocin. Positive clones were selected and the expression of ISG20 was evaluated in the cytosol (endogenous) and supernatant (secreted) by western blot analysis (Fig.6C). Positive clones were selected based on the intracellular expression of ISG20, but more importantly, the detection of ISG20 expression in the supernatant; confirming the secretion of the protein (Fig.6C). No detectable ISG20 protein was found in the supernatant or cell lysate of the negative clone or non-transfected CHO cells (Fig.6C).

**Figure 6.**
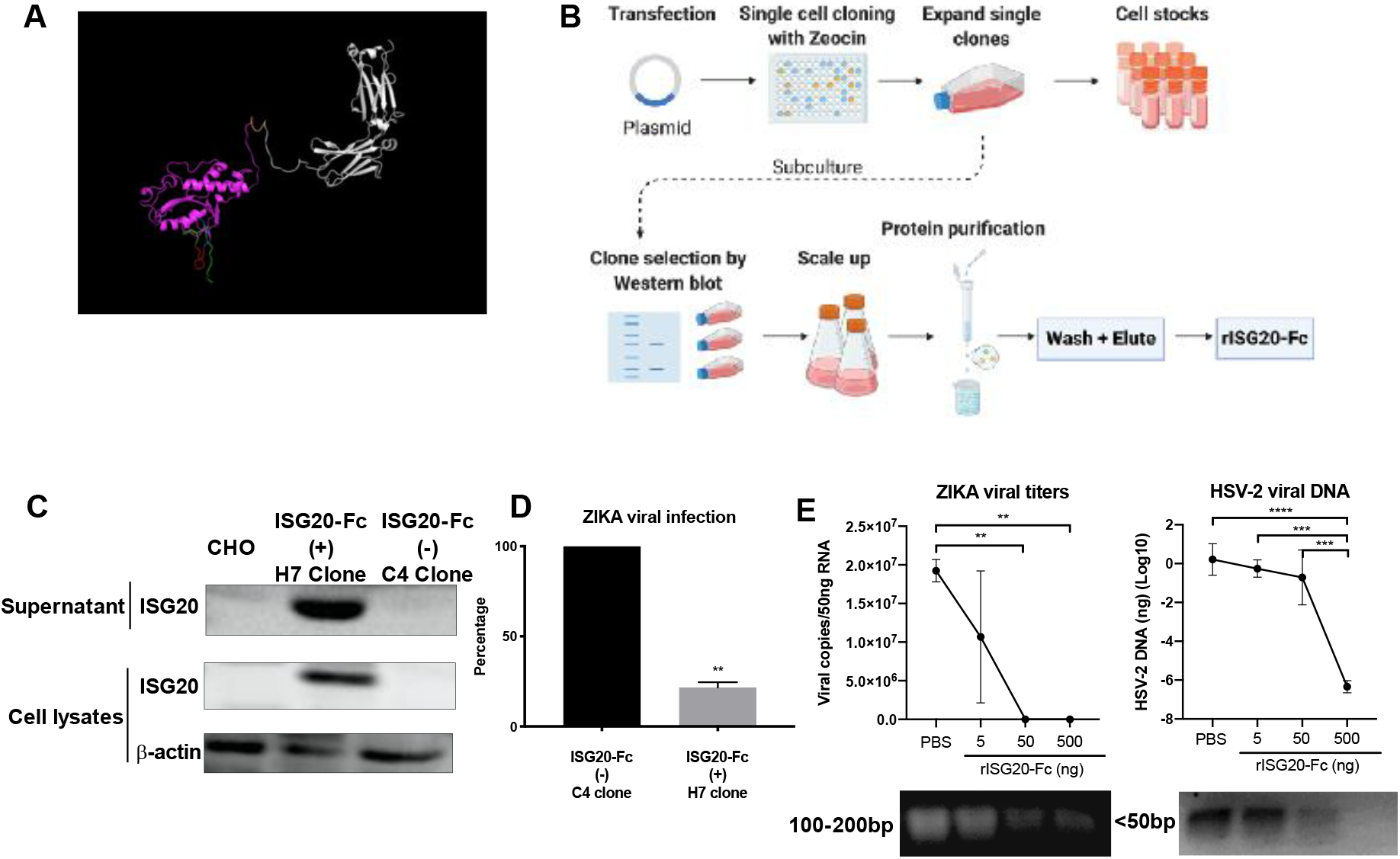
Design and characterization of the anti-viral effect of recombinant ISG20 protein. **(A)** Design of the recombinant ISG20-Fc protein. The components are: 1) ISG20 sequence (Pink); 2) Linker (Orange); 3) Fc domain (White); 4) Tev sequence, cleavage between Q and G (Cyan); 5) Signal sequence (ENPP7) (Red); 6) His Tag (Green). **(B)** Workflow of clone production and selection. **(C)** Selection and verification of positive clones. In the positive clone, ISG20 protein expressed both in the cytosol (endogenous) and supernatant (secreted) compared to the non-transfected CHO cells and transfected negative clones. CHO, Chinese hamster ovary cell without transfection; β-actin served as a loading control. **(D)** ISG20 secretion in the positive clone supernatant significantly decreased ZIKV infection in trophoblast cells. Supernatants from negative and positive clones were collected and added to ISG20^−/−^ Sw.71 trophoblast cells together with ZIKA virus (MOI=2) for 1 h, followed by refreshing with new growth media for 48 h. RNA was then collected to determine viral titers by qRT-PCR. Data represent as mean ± SEM from two independent experiments. \*\**p* < 0.01. **(E)** Recombinant ISG20-Fc degrades ZIKA viral RNA and HSV2 viral DNA. 50ng purified viral RNA (ZIKV) or DNA (HSV-2) were incubated with increasing concentrations of rISG20-Fc (5, 50, 500 ng) in the presence of RNase inhibitor for 90min at 37°C followed by quantification of viral titers by qRT-PCR, and agarose gel was used to evaluate the RNA degradation by electrophoresis. The representative picture of agarose gel is presented. rISG20-Fc was able to degrade both ZIKV RNA and HSV-2 DNA in a dose dependent manner; however, rISG20-Fc was more efficient in degrading viral RNA than DNA. Data represent as mean ± SEM from three independent experiments. \*\**p* < 0.01. \*\*\**p* < 0.001 and \*\*\*\**p* < 0.0001.

Next, we collected the conditioned media from the positive (H7) and negative (C4) clones and added that to the ISG20^−/−^ Sw.71 trophoblast cells together with ZIKA virus (MOI=2) for 1 h, followed by refreshing with new growth media for 48 h. RNA was collected to determine viral titers by qRT-PCR. The conditioned media from the positive clone H7, significantly reduced ZIKV viral titers compared to the negative clone C4 (Fig.6D). This data suggests that the secreted form of ISG20 preserves its RNase activity and shows anti-ZIKA effect. Consequently, we proceeded to purify the recombinant ISG20-Fc protein for further characterization (see Material & Methods).

### 7. Anti-viral activity of recombinant ISG20-Fc protein

To test if the exonuclease activity of our recombinant ISG20-Fc (rISG20-Fc) was preserved, we first examined its ability to degrade viral RNA or DNA. We used two different viruses as the substrates for this experiment: RNA virus (ZIKA) and DNA virus (HSV-2). 50ng purified ZIKV RNA or HSV-2 DNA were incubated with increasing concentrations of rISG20-Fc (5, 50, 500 ng) for 90min at 37°C in the presence of RNase inhibitor to exclude the effect of exogenous RNase, followed by quantification of viral titers by qRT-PCR. rISG20-Fc was able to degrade both ZIKV RNA and HSV-2 DNA in a dose dependent manner, however, rISG20-Fc was more efficient in degrading viral RNA than DNA (Fig.6E).

### 8. In vivo efficacy of rISG20-Fc inhibiting ZIKA viral replication in IFNAR1^−/−^ pregnant mice

Having shown that rISG20-Fc maintained its exonuclease activity, we then tested whether rISG20-Fc could have an anti-viral effect by using the IFNAR1^−/−^ mice, which are highly sensitive to ZIKV infection (Lazear et al., 2016). Adult (8-12 weeks of age) IFNAR1^−/−^ pregnant mice were intraperonteially (i.p.) infected with 1×10^5^ pfu ZIKV or 1% FBS DMEM/F12 media (vehicle) at day E8.5 (Fig.7A). Afterwards, animals received three doses of rISG20-Fc (1mg/kg) i.p. 1 h post ZIKV infection and on days E9.5 and E10.5 of their pregnancy. Control animals were injected with PBS at the same time points (Fig.7A). On E14.5, mice were sacrificed and organs were collected to assess viral titer. In control mice, macroscopic evaluation of the uterus showed the presence of fetal death and resorptions in ZIKV infected mice (Fig. 7B). However, rISG20-Fc treatment rescued this phenotype by decreasing the number of resorptions and preventing fetal death (Fig.7B and Sup.Fig.5A). No differences in the number of implantation sites were found between the two groups, which confirms the protective effect of rISG20-Fc (Sup.Fig.5B).

**Figure 7.**
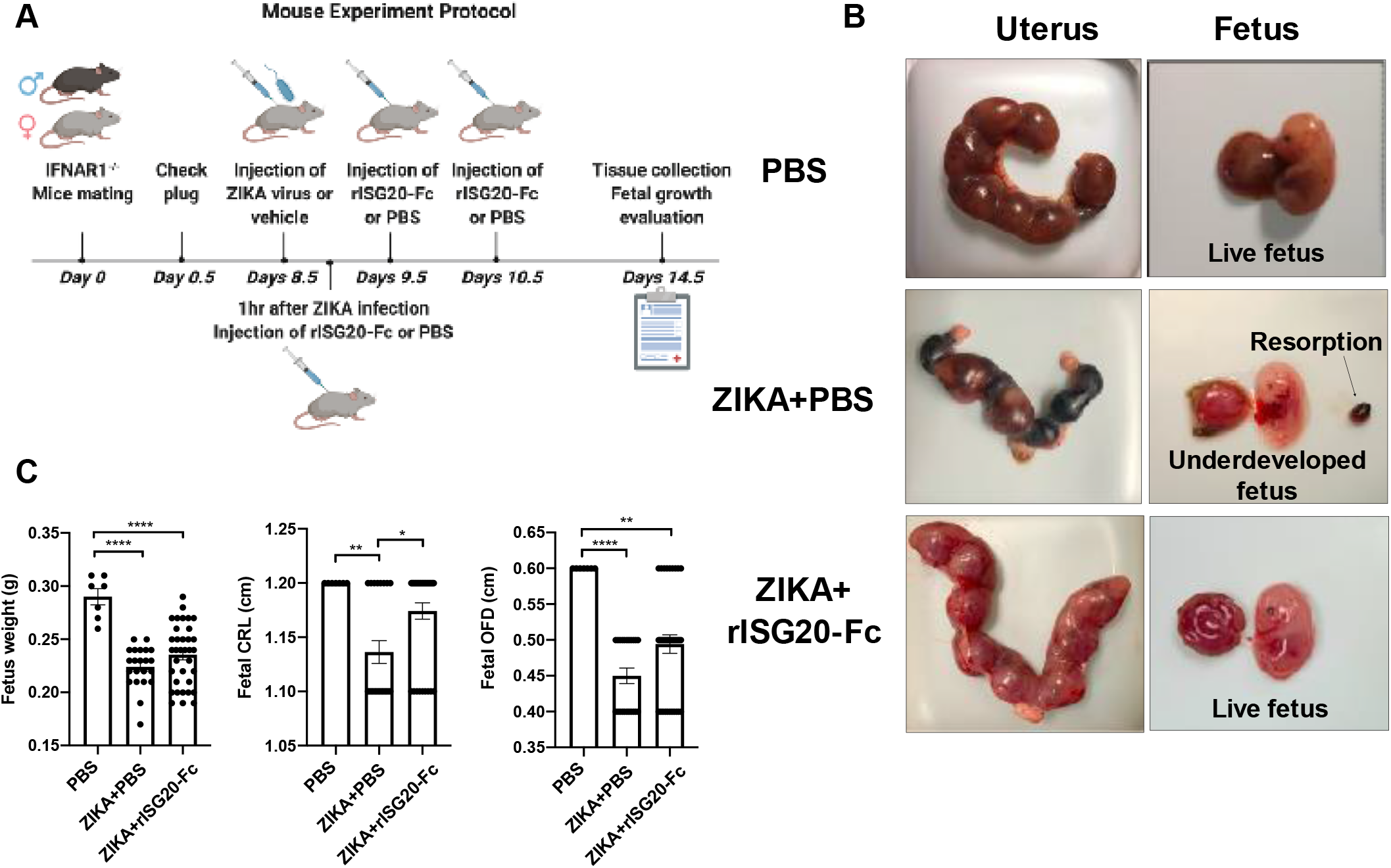
In vivo efficacy of rISG20-Fc improving pregnancy outcome in IFNAR1^−/−^ pregnant mice during ZIKV infection. **(A)** Mouse experiment protocol. Adult (8-12 weeks of age) IFNAR1^−/−^ pregnant mice were infected i.p. with 1*10^5^ pfu ZIKV or 1% FBS DMEM/F12 media (vehicle) on embryo day 8.5 (E8.5). 1 h after ZIKV/vehicle injection, treatment with rISG20-Fc (1mg/kg) or PBS (control) was administrated i.p. to the pregnant mice. On E9.5 and E10.5, same protein/PBS injection was performed on the pregnant mice. On E14.5, the mice were sacrificed and organs were collected for viral titer quantification. N=2 for control PBS group, and n=3-4 for treatment groups. **(B)** Macroscopic evaluation of the pregnant uterus and fetus. Compare to the fetal death and reabsorptions in ZIKV-infected mice, rISG20-Fc treatment rescued this phenotype by preventing fetal death and reabsorptions. **(C)** rISG20-Fc treatment improves fetal development. Measurement of fetal weight, crown-rump length (CRL) and occipitofrontal diameter (OFD) were evaluated when sacrificing mice on E14.5. ZIKV infection significantly inhibited fetal development, while rISG20-Fc treatment could significantly increase CRL. \**p* < 0.05, \*\**p* < 0.01 and \*\*\*\**p* < 0.0001.

Next, we analyzed the impact of ZIKV infection and treatment with rISG20-Fc on fetal development. As previously reported (Uraki et al., 2017), ZIKV infection has major negative impacts on fetal development as demonstrated by a significant decrease in fetal weight, crown-rump length (CRL), and occipitofrontal diameter (OFD) (Fig.7C). Treatment of ZIKV-infected pregnant mice with rISG20-Fc significantly improved fetal CRL when compared to the infected group, which suggests that rISG20-Fc can promote fetus development in the uterus (Fig.7C).

We then evaluated the anti-viral effect of rISG20-Fc treatment by quantifying ZIKV titers on the maternal and fetal side. On the maternal side, we observed a significant decrease in ZIKV titers in the maternal serum and spleen in mice that were treated with rISG20-Fc (Fig.8A). On the fetal side, we saw a significant decrease in ZIKV titers in the fetal brain of rISG20-Fc treated mice (Fig.8B). Interestingly, there was no observed difference in ZIKV titer in the placenta between the two groups. This suggests that rISG20-Fc can block viral transmission from the placenta to fetus, thus protecting the fetus from the detrimental effects of ZIKV infection.

**Figure 8.**
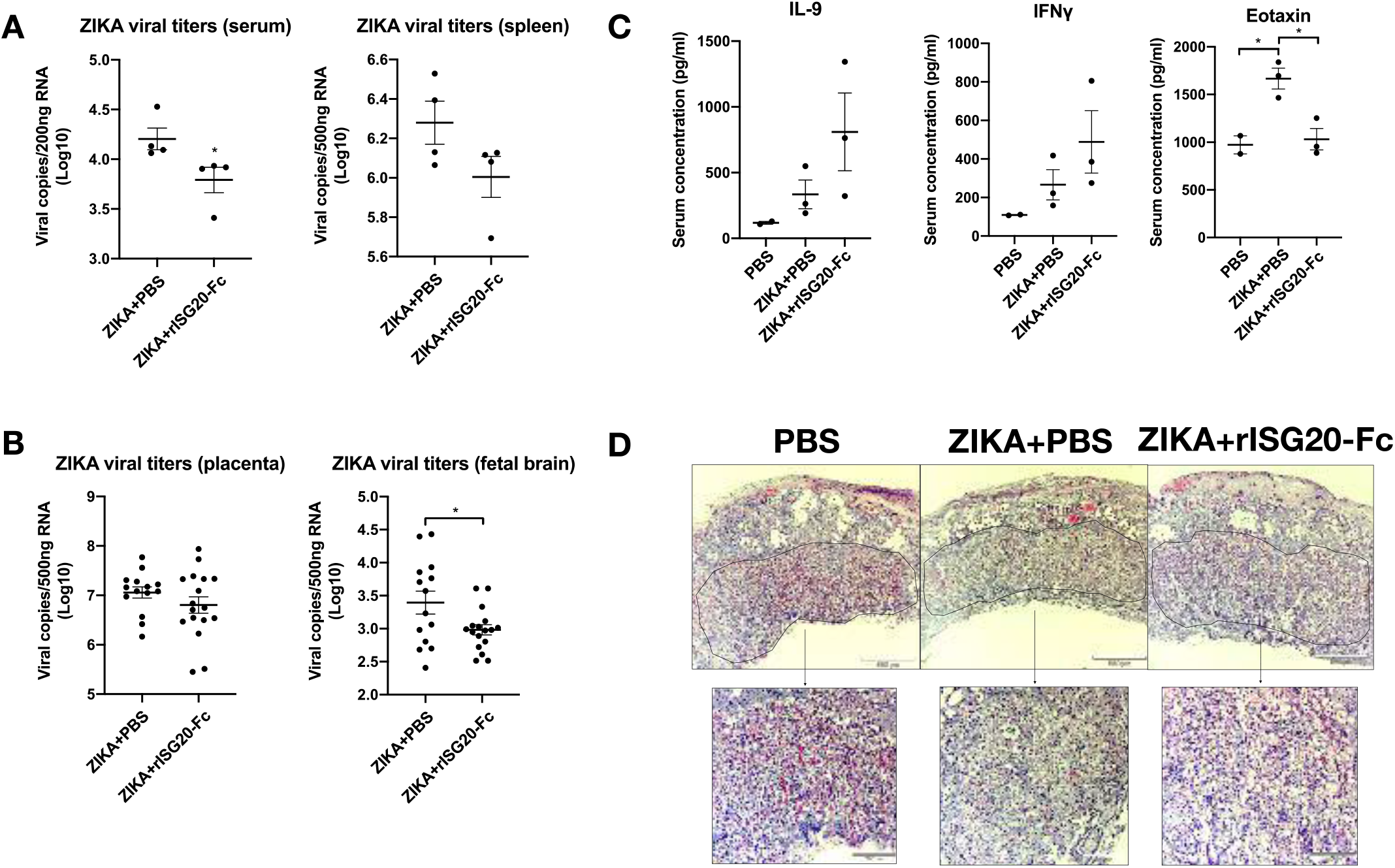
In vivo efficacy of rISG20-Fc inhibiting ZIKA viral replication and promoting placental integrity in IFNAR1^−/−^ pregnant mice. **(A)** rISG20-Fc treatment alleviates maternal viral burden. Note that rISG20-Fc treatment decreased the maternal serum and spleen viral titers. **(B)** rISG20-Fc treatment reduces ZIKV titers in the fetal brain. Although there was no difference of ZIKV titer in the placenta, at the fetal side, there was a significant decrease of viral titer in the fetal brain, suggesting rISG20-Fc can block viral transmission from placenta to fetus. N=4 for each group, and 3-4 placentas/fetal brains from every mouse were analyzed. \**p* < 0.05. **(C)** rISG20-Fc treatment increases the expression of IL-9, IFNλ, and decreases Eotaxin in maternal circulation. N=2 for PBS control group, and n=3 for every treatment group. \**p* < 0.05. **(D)** Representative hematoxylin and eosin staining of the mouse placenta on E14.5. Labyrinth layers were marked with a solid line. Note the major alteration on placenta structure characterized by multifocal loss of tissue architecture (necrosis) and the defective blood vessel formation of labyrinth in ZIKV+PBS group, and rISG20-Fc treatment improves vascularity and decreases decidual edema and cellular fragmentation at the labyrinth.

### 9. Impact of rISG20-Fc treatment on placental integrity

Finally, we explored if treatment with rISG20-Fc could modulate the immunological response elicited by ZIKV infection on the maternal and fetal side. Thus, we evaluated maternal serum cytokine expression of ZIKV infected dams treated with rISG20-Fc or vehicle control. Interestingly, we observed that rISG20-Fc treatment had a modulatory effect on the anti-viral response by enhancing the expression of IL-9 and IFNγ (potent anti-viral cytokines) and reducing inflammatory cytokines (Endotaxin) (Fig.8C).

Next, histologic analysis of placenta samples from each group was performed to determine the impact of the ZIKV-induced inflammatory process on the placenta and if this could be countered with rISG20-Fc treatment. Thus, uteroplacental units were collected at E14.5 (day 6 post infection), and H&E staining was performed. All histological samples were analyzed in a blinded manner by an independent animal pathologist. As shown in figure 8D, placentas obtained from animals infected with ZIKV showed major alteration on placenta structure characterized by multifocal loss of tissue architecture (necrosis) in the labyrinth, apparent damage of the vascularity evidenced by decrease blood vessel density, edema were observed only in the decidua of infected mice, necrosis and inflammation foci were observed in the labyrinth of infected mice. Additional pathologic changes present within the labyrinth of infected mice included an overall tissue hypereosinophilia, nuclear pyknosis, and cellular fragmentation (Fig.8D). Treatment with rISG20-Fc reversed some of these changes, including increased vascularity, decreased decidual edema, and cellular fragmentation at the labyrinth (Fig.8D). This data suggests that rISG20-Fc treatment can contribute to placenta integrity during ZIKV infection and facilitate fetal development.

## Discussion

We report for the first time, the characterization of an effective IFNβ-mediated anti-ZIKV response in trophoblast cells and the identification of ISG20, an endogenous anti-viral protein, which is naturally produced by the placenta/trophoblast, as an earlier anti-viral exonuclease that is critical for stopping the viral cycle by promoting viral RNA degradation and providing immune regulation. Furthermore, we successfully established a recombinant ISG20-Fc protein that maintains its exonuclease activity and effectively decrease viral titers in vitro and in vivo.

ZIKV infection during pregnancy remains a significant concern due to the major impacts on fetal development, such as microcephaly and other catastrophic fetal malformations (Alvarado and Schwartz, 2017, Schwartz, 2017a). Due to the high sensitivity of the infection during pregnancy, ZIKV is now included in the list of viral infections dangerous for pregnancy known as “TORCH” pathogens (Coyne and Lazear, 2016, Schwartz, 2017b), along with *Toxoplasma gondii*, rubella virus, cytomegalovirus (CMV) and herpes simplex virus (HSV), which are considered as the major causes of fetal morbidity and mortality. Unfortunately, there is no effective vaccine or treatment for ZIKV infection. Therefore, development of new therapies is needed.

In this study, we characterized the natural anti-viral response to ZIKV infection and identified early responders that are important for stopping viral replication. The host innate immune response to viral infection is critical for controlling viral replication through the induction of type I IFNs and ISG expression (Schneider et al., 2014, Ivashkiv and Donlin, 2014, Espert et al., 2003). ISGs are considered as the anti-viral effectors of the type I IFN response(Gifford et al., 2007). These effectors may function by targeting different stages of viral life cycle, such as inhibition of virus entry, viral RNA replication, viral particle assembly, and release (Schneider et al., 2014, Sadler and Williams, 2008, Munakata et al., 2008, Jiang et al., 2008). Hence, with the absence of type I IFN-ISGs signals, virus will be likely to replicate more and destroy the tissues. For example, in ZIKV infection mouse model, defects in the type I IFN signaling axis such as the lack of IFNAR1 (IFNAR1^−/−^) or IRF3/5/7 have been shown to increase the sensitivity of ZIKV burden in multiple organs, such as the brain, spinal cord and testes. In these same models, ZIKV-infected IFNAR1^−/−^ mice developed neurological diseases and succumbed to ZIKV infection (Lazear et al., 2016). Moreover, ZIKV infection in IFNAR1^−/−^ mice during early pregnancy resulted in fetal demise, which was associated with the infection of placenta and fetal brain (Miner et al., 2016b).

In the placenta, trophoblast cells are able to mount a strong type I IFNβ response, which is characterized by the expression of multiple ISGs (Kwon et al., 2018). We found here a significant increase in IFNβ transcription and translation during ZIKV infection in the Sw.71 first-trimester trophoblast cell line and primary trophoblast cell cultures. This was then followed by the expression of a number of critical anti-viral ISGs involved in this response, including *Isg20, Mx1, Oas1, Isg15, CH25H, Trim22, Tetherin* and *Viperin*. Among these ISGs, ISG20, which is a 3’-5’ exonuclease, stands out for its early disruptive role in viral RNA replication. Prior studies have noted the importance of the anti-viral effect of ISG20 in several viruses, including Hepatitis B virus (HBV), Hepatitis C virus (HCV), West Nile virus, Dengue virus, and Human Immunodeficiency virus (HIV) (Espert et al., 2005, Jiang et al., 2008, Liu et al., 2017), and confirmed that ISG20-mediated antiviral activity mainly depends on its exonuclease activity (Zheng et al., 2017).

Therefore, we further focused on characterizing ISG20 expression and function in trophoblast cells and observed that IFNβ and Poly (I:C) are both strong inducers of ISG20 expression. Poly (I:C) is a synthetic double-stranded RNA (dsRNA), which mimics a viral infection and activates the TLR3 and cytoplasmic dsRNA sensors signaling pathways leading to increase of IFNβ expression (Kato et al., 2006, Watanabe et al., 2011). Exposure of trophoblast cells to Poly (I:C) induces IFNβ and ISG20 expression. This further supports the findings that trophoblast cells are capable of mounting an anti-viral response, which is characterized by the expression of IFNβ and ISG20. Noteworthy, we did not observe ISG20 protein expression in the cells during regular culture conditions although we did detect high levels of mRNA expression. Only upon viral infection or IFNβ treatment can we transiently detect the expression of ISG20. The regulatory pathways associated with this quick and transient expression is under further investigation.

To determine the role of ISG20 in the anti-ZIKV response, we established a trophoblast cell line where ISG20 was knocked out. We found that without ISG20, activation of the IFNβ pathway and induction of downstream anti-viral ISGs was compromised, which suggest a regulatory role for ISG20 in the expression and function of other ISGs associated with type I IFN response. Importantly, lack of ISG20 weakens the protective effect provided by IFNβ against ZIKV infection, which further confirms the central role of ISG20 in the IFN-β mediated response against ZIKV infection. Our findings are in line with a previous report showing that ISG20 exerts antiviral activity through upregulation of other IFNβ response proteins (Weiss et al., 2018).

In order to repurpose this anti-viral gene as a therapeutic for ZIKV infection, we designed a recombinant ISG20 protein. To design the ISG20 protein, we added an Fc domain of human neonatal IgG1 that functions to increase protein stability and facilitate the uptake by cells(Czajkowsky et al., 2012, Bell et al., 2013). Since multiple cell types express the Fc receptor, including immune cells, mucosal epithelial cells, and trophoblast cells (Lozano et al., 2018), it is plausible that our recombinant ISG20-Fc may bind to Fc receptors on the surface of cells to facilitate endocytosis. More importantly, it is well established that IgG transfer from the mother to fetus across the placenta is mediated by neonatal Fc receptors expressed by the placenta (Lozano et al., 2018). Consequently, the presence of the Fc domain in rISG20-Fc may allow the binding of the recombinant protein to trophoblast cells and facilitate its transport from the maternal circulation into the fetal tissues. Furthermore, we thoroughly examined the exonuclease activity of rISG20-Fc by incubating the recombinant protein with purified ZIKV RNA and HSV-2 DNA. rISG20-Fc is highly effective at degrading ZIKV RNA as well as HSV-2 DNA. However, as previously reported (Moser et al., 1997, Nguyen et al., 2001), rISG20-Fc displays a higher efficacy towards the degradation of viral RNA rather than DNA.

The placenta is an extremely important immune organ, which acts as a physical and immunological barrier to block the transmission of infectious agents to fetus during pregnancy(Mor et al., 2017, Mor and Cardenas, 2010). Due to the invasive nature of the placenta in both humans and, more limited, in mice, trophoblasts make direct contact with maternal circulation. Previous studies have shown that ZIKV infects human trophoblasts (Bayer et al., 2016, Lazear et al., 2016, Quicke et al., 2016), and the primitive trophoblasts formed in the early stage of pregnancy are more sensitive to ZIKV infection compared to the later stage after the formation of syncytium (Sheridan et al., 2017). Therefore, we postulated that use of rISG20-Fc could be an important tool to provide a protective effect in the developing placenta during early pregnancy. Indeed, when tested in vivo using the IFNAR1^−/−^ mouse model, rISG20-Fc was found to be effective in protecting the fetus from ZIKV infection. While IFNAR1^−/−^ pregnant mice infected with ZIKV showed a high number of resorptions and underdeveloped fetuses; treatment with rISG20-Fc was able to rescue the fetus and improve their growth by preventing ZIKV-induced placental damage.

Furthermore, it is likely that rISG20-Fc has systemic immune modulatory function as it can also decrease the viral titer in maternal circulation by facilitating the secretion of anti-viral cytokines, such as IL-9 (Yu et al., 2016) and IFNλ (Kang et al., 2018); in addition to balancing the overexpression of pro-inflammatory factors, such as Eotaxin (Rankin et al., 2000). Unexpectedly, we observed similar levels of ZIKV titers in placental samples from treated and control groups. No observed decrease of viral titers in the placenta suggests that rISG20-Fc may be transferred from maternal circulation to the fetus through the Fc receptor in placenta and then degrades viral RNA on the fetal side, as demonstrated in figure 9.

**Figure 9.**
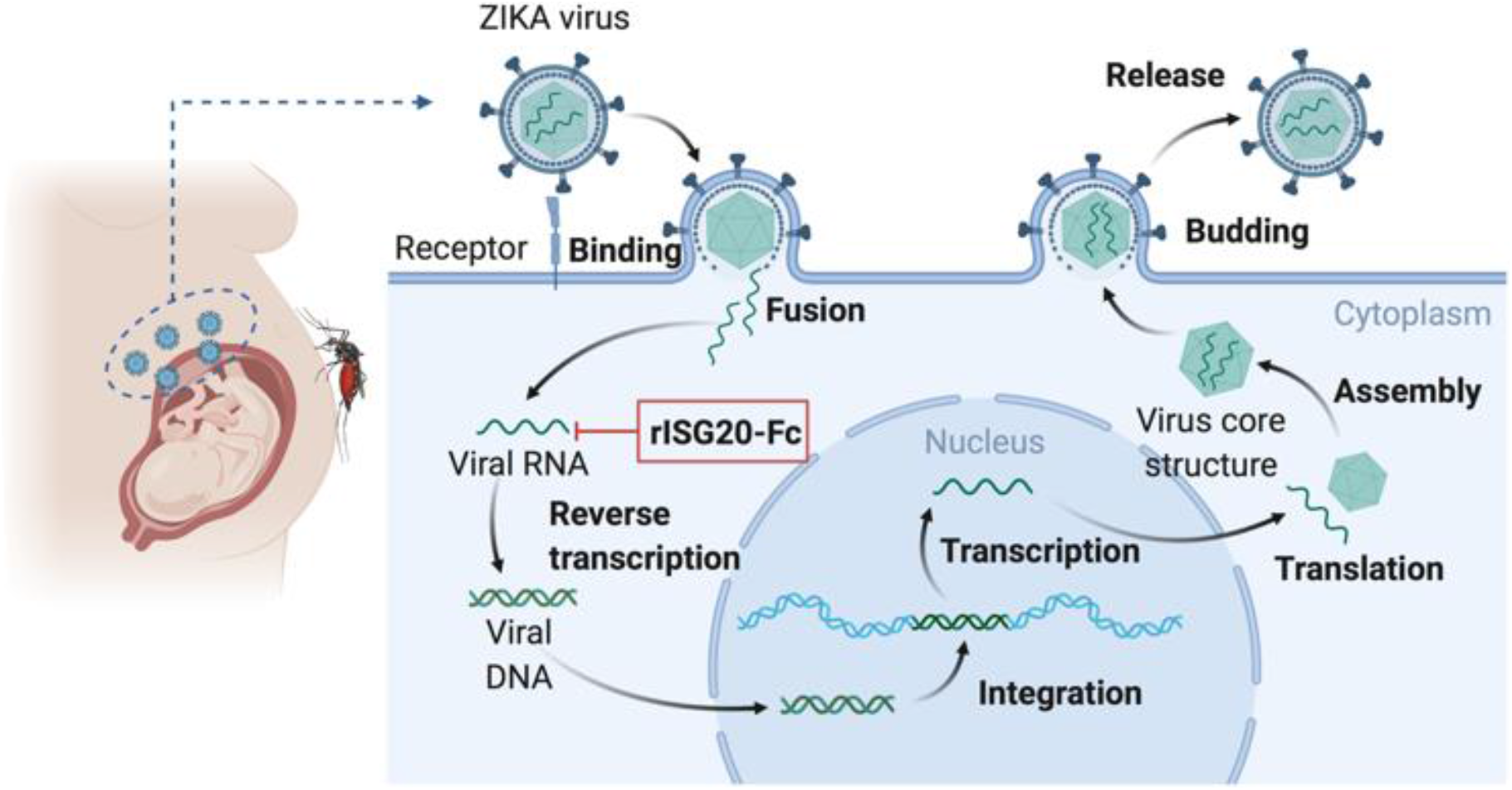
The diagram of rISG20-Fc inhibiting ZIKA viral replication and decreases viral infection in fetus. When ZIKA virus infects the pregnant woman, the virus will reach the placenta and transmit to the fetus. In the ZIKV replication cycle, it starts with the virus binding to host cell surface receptors, leading to endocytosis of the virus. Internalized viral particles release the viral RNA into the cytoplasm of the host cell and start replication. During its replication, rISG20-Fc degrades ZIKV RNA and inhibits the subsequent transcription and translation. Therefore, less new virus particles are released from the placenta to fetus.

Altogether, we identified ISG20 as a central effector in the IFNβ-induced response to ZIKV infection in trophoblast cells. We have successfully generated a recombinant form of ISG20 that maintains its exonuclease activity and has protective effect on an animal model that is sensitive to ZIKV infection. We show that rISG20-Fc possess a strong anti-viral exonuclease activity in addition to immune modulatory functions. Thus, it is plausible that our recombinant protein can be applied towards other types of viral infections, especially for RNA virus infection. The findings from this study provide a new insight on the pathways regulating ZIKV infection and offer a valuable tool to combat ZIKV infection during pregnancy.

## Material and Methods

### Cell culture and infection

Immortalized human trophoblast Sw.71 cells and human endometrial stroma cells (HESC) were cultured in DMEM/ F12 or DMEM supplemented with 10% FBS, 10 mm HEPES, 0.1 mm MEM non-essential amino acids, 1 mm sodium pyruvate, and 100 U/ml penicillin/streptomycin (Life Technologies; Waltham, MA, USA) under 5% CO_2_ at 37°C. For viral infections, cells were seeded in 6-well plates at 1.75×10^5^ cells per well, the next day, ZIKV was added to the cells for 1 h incubation (at indicated MOI) with gentle agitation every 20 min, after 1 h, the inoculum was removed and the cells were washed twice with phosphate-buffered saline (PBS), then cells were maintained in 10% FBS DMEM/F12 media for the duration of the experiment. At indicated time points after infection, cell pellets and conditioned media were collected for downstream analysis.

### Human primary trophoblast isolation and culture

Human primary trophoblast cells were isolated from first trimester elective terminations as previously described (Straszewski-Chavez et al., 2009). A signed written consent form was obtained from the patients. The use of placental tissues, specimens and consent forms was approved by the Yale University Human Investigation Committee (#2000021607). The tissue specimen was collected in cold, sterile phosphate-buffered saline (PBS) and immediately transported to the laboratory for cell culture preparation. Briefly, first trimester placental villous tissues were cut and digested in PBS supplemented with 0.25% Trypsin (Gibco, Grand Island, NY, USA) for 10 min at 37°C with gentle agitation. An equal volume of 10% FBS (Gibco, Grand Island, NY, USA) and Dulbecco’s Modified Eagle Medium (DMEM) (Gibco, Grand Island, NY, USA) was added to inactivate the trypsin. The supernatant was collected and centrifuged at 1500 rpm at room temperature for 10 min. The pellet was resuspended in 5 ml DMEM media supplemented with 10% FBS. This suspension was laid over lymphocyte separation media (ICN Biomedicals, Inc., Aurora, OH, USA) and centrifuged at 2000 rpm for 20 min. The interface containing the trophoblast cells was collected and centrifuged at 1500 rpm for 10 min. Cells were resuspended in DMEM with 10% FBS and then plated on a 6-well plate to grow.

### Knockout of ISG20 using CRISPR/Cas9

ISG20 was knocked out in Sw.71 first-trimester trophoblast cell line using CRISPR/Cas9. The guide RNA for ISG20 was designed using the CRISPR design tool from the Zhang Laboratory at MIT (CRISPR.mit.edu) (Sanjana et al., 2014, Shalem et al., 2014). Two DNA oligos were synthesized as follows: Sense: 5’-CACCGCAGCACCGTGGACGTTCACG-3’, Antisense: 5’-AAACCGTGAACGTCCACGGTGCTGC-3’. Oligos were phosphorylated using T4 polynucleotide kinase and annealed by heating equimolar amounts to 95°C and cooling slowly to room temperature. This resulting guide was introduced to lentiCRISPRv2GFP plasmid using BsmBI restriction sites, and lentiCRISPRv2GFP was a gift from David Feldser (Addgene plasmid # 82416) (Walter et al., 2017). 10 μg of the resulting plasmid was co-transfected with 8 μg of packaging plasmid pCMV-VSV-G and 4 μg of envelope plasmid psPAX2 in the presence of 60 μg of polyethylenimine (PEI) into HEK293T cells in a 100-mm dish. pCMV-VSV-G was a gift from Bob Weinberg (Addgene plasmid # 8454) (Stewart et al., 2003), and psPAX2 was a gift from Didier Trono (Addgene plasmid # 12260). Then packaged viral particles were collected by ultracentrifugation and were transduced into the Sw.71 cells. The cells were sorted based on the GFP signal by fluorescence activated cell sorting (FACS) following transduction. The deletion of ISG20 was confirmed using Sanger sequencing technique performed by GENEWIZ, and the overall efficiency was 91.1% analyzed by TIDE (Tracking of Indels by Decomposition). Moreover, we verified the protein expression by Western blot (Sup. Fig. 4A).

### Virus

ZIKV strain FSS 13025, which was originally isolated in Cambodia in 2010, was obtained from the World Reference Center for Emerging Viruses and Arboviruses at University of Texas Medical Branch, Galveston as previously described (Aldo et al., 2016b). ZIKA virus was propagated in African green monkey kidney (Vero) cells by infecting the monolayer with viral stock. When the cytopathic effect was observed in the whole monolayer, the infected supernatant was collected and centrifuged. The virus stocks were aliquoted and stored at −80°C. The viral titer of viral stock was determined by plaque assay.

### Plaque assay

The infectivity of the virus was determined by plaque assay in Vero cells. Briefly, Vero cells were plated into 12-well plates at 2.5×10^5^ cells/well and inoculated with 200ul of 10-fold serial dilutions of viral stocks and incubated at 37°C for 1 h with gentle agitation. After inoculation, Vero cells were overlaid with media containing DMEM (Gibco, #11965-084), 2% FBS, and 0.6% Avicel (FMC, # CL-611). Cells were maintained at 37°C in 5% CO_2_ for 5 days. After 5 days incubation, overlays were aspirated and the cells were fixed in 4% formaldehyde solution in PBS before staining with 1% crystal violet in 20% methanol. Viral plaques were photographed and each plaque was counted as a plaque-forming unit (PFU). Viral titer was calculated as PFU/ [volume virus (mL) × (dilution factor)].

### Production and purification of recombinant ISG20-Fc protein

For the production of recombinant ISG20 protein from mammalian cell cultures, we designed the plasmid and transfected it into Chinese Hamster ovary cells (CHO). Briefly, an artificial gene sequence was obtained from Life Technologies, which coded for the following: 5’ BamHI restriction site, cell export signal sequence from hENPP7, 10xHis tag for nickel purification, Tev sequence (to enable cleavage of the protein), full-length human ISG20, a linker (Gly-Ser-Gly-Ser-Gly), the Fc domain of human IgG1, and a 3’ EcoRI restriction site. In order to increase the binding affinity of our fusion protein with the Fc receptor, we changed three amino acids within the Fc region by using the QuikChange II site directed mutagenesis kits (Agilent, #200523), introducing M257Y, S259T, and T261E mutations simultaneously in the Fc region.

Mutagenesis primers are as follows: MST Fwd: 5’-ccccaaagcccaaagacactctgtatatcaccagggagcctgaagttacatgcgtcgttgt-3’ MST Rev: 5’-acaacgacgcatgtaacttcaggctccctggtgatatacagagtgtctttgggctttgggg-3’ All of this sequence had been codon optimized for efficient expression in CHO cells. Using BamHI/EcoRI, this cassette was subcloned into pcDNA4 (Invitrogen, #V102020) vector, because the carrier vector in which the cassette was supplied lacked a promoter. Once completed, the plasmid was transfected into CHO cells using polyethyleneimine, and cells were then selected for plasmid integration with Zeocin treatment.

After transfection and Zeocin selection, the cells were dissociated and serial diluted into a selection media containing 150 μg/ml Zeocin for 2 weeks to establish stable single cell clones. After the expanding of single clones, positive clones were selected by evaluating the protein expression of ISG20-Fc in the cytosol (endogenous) and supernatant (secreted) by western blot analysis. Additionally, conditioned media from stable cell clones was used to assess anti-ZIKA activity in vitro, and the clones with the highest efficacy were expanded and adapted for suspension growth in Pepro AF-CHO serum free media (Peprotech). Six liters of media at 5×10^6^ cells/ml was centrifuged at 1000xg for 30 minutes and the secreted protein was purified to homogeneity as previously described (Albright et al., 2015) using an ÄKTA Pure 25M equipped for multi-step automated purification with modifications. First, many non-specific CHO cell secreted proteins were eliminated from the active fraction by ammonium sulfate precipitating, final concentration of 23% (NH_4_)_2_SO_4_. The remaining soluble protein was further purified by sequential nickel column and MabSelect prismA affinity column purification followed by an S75 sizing column. Purified proteins were stored as frozen stocks in PBS at −80°C.

### RNA degradation assay

Purified ZIKV RNA and HSV-2 DNA were extracted using QIAamp MinElute Virus Spin Kit (Qiagen, catalog no. 57704), and 50ng purified viral RNA/DNA was incubated with the recombinant protein at different concentrations for 90min at 37°C in the presence of SUPERase In™ RNase inhibitor (Invitrogen, catalog no. AM2694), and the resulting RNA/DNA was tested for viral copies by qRT-PCR. Subsequently, the PCR product was used for agarose gel electrophoresis to evaluate the degradation of RNA/DNA as described (Aranda et al., 2012).

### Mouse experiments

The IFNAR1−/− (B6.129S2-Ifnar1tm1Agt/Mmjax) mice were obtained from the Jackson Laboratory (Bar Harbor, ME) and bred in a specific-pathogen-free facility at Wayne State University. Adult (8-12 weeks of age) IFNAR1^−/−^ mice were set up for timed-mating and the plug day was considered as embryonic days E0.5. On embryonic days E8.5, the plug-positive mice were inoculated intraperitoneally (i.p.) with either 1×10^5^ plaque-forming units (pfu) of ZIKA virus (in 100 μL volume) or 1% FBS DMEM/F12 media (vehicle). 1 h after ZIKV/vehicle injection, treatment with rISG20-Fc (1mg/kg) or PBS (control) was administrated i.p. to the pregnant mice. On E9.5 and E10.5, same protein/PBS injection was performed on the pregnant mice. On E14.5, the mice were sacrificed for tissue collection.

Pregnancy outcome parameters including the number of implantations and resorptions were recorded, and fetus development was evaluated by comparing the weight, fetal crown-rump length (CRL) and occipitofrontal diameter (OFD) between groups. Maternal spleen, placentas, fetal brain were collected and stored in RNA*later*™ stabilization solution (Invitrogen, catalog no. AM7021) in −80°C for viral titer quantification. Furthermore, maternal serum was used for cytokine expression analysis by Luminex (BioRad) and some placentas from each group were stored in 4% paraformaldehyde for haematoxylin and eosin staining.

### RNA extraction and Real-time PCR analysis

Cells were collected and total RNA was extracted using the RNeasy Mini kit (Qiagen, catalog no. 74106) according to the manufacturer’s instructions. Viral RNA from mice serum was extracted using QIAamp MinElute Virus Spin Kit (Qiagen, catalog no. 57704) according to the manufacturer’s protocol. RNA from mice tissue was extracted as described. Briefly, tissue was homogenized in Trizol (Ambion; Waltham, MA, USA) in 1.0-mm zirconium beads (Benchmark; Sayreville, NJ, USA) for two cycles at 400 *g* for 2 min. Supernatant was then transferred to a new tube and incubated for 5 min at room temperature (RT). 200 μl chloroform/ml of Trizol was added, shaken for 15 s, and then incubated for 3 min at RT. Samples were centrifuged at 12,000 *g* for 15 minutes at 4°C, and aqueous phase was transferred to new tubes. Then, 500 μl of 100% isopropanol was added, incubated at RT for 10 min, and centrifuged at 12,000 *g* for 10 minutes at 4°C. Supernatant was then discarded, and cell pellet was washed twice with 500 μl of 100% ethanol and centrifuged at 12,000 *g* for 5 min. The pellet was then washed twice with 1 ml of 75% ethanol, vortexed briefly, then centrifuged at 7,500 *g* for 5 min at 4°C, and then air-dried for 10 minutes. The pellet was resuspended in 50 μl of RNase-free water.

RNA concentration and purity were assessed using spectrophotometric analyses of 260/280 ratios, and only samples with values of 1.8 or higher were used for PCR analysis. One microgram of RNA was reverse-transcribed for each sample using Bio-Rad (Hercules, CA, USA) iScript cDNA synthesis kit. iTaq Universal SYBR Green Supermix (BioRad) and gene-specific primers (Sequences listed in Supplemental Table 1) were added to the RT reactions that were diluted 1:5 with nuclease-free water and run on the CFX96, C1000 system qPCR machine (BioRad). Values were normalized to GAPDH and calculated with 2^−ΔΔCt^ method as described (Livak and Schmittgen, 2001).

### Virus titer quantification by qRT-PCR

ZIKA viral titer was quantified by one-step quantitative reverse transcriptase PCR (qRT-PCR). ZIKA Viral RNA was isolated from Zika virus stock using QIAamp MinElute Virus Spin Kit (Qiagen, catalog no. 57704) to create a standard curve using serial 10-fold dilutions of ZIKV RNA. One microgram of RNA from samples was run on CFX96, C1000 system qPCR machine (BioRad) using a one-step PCR mix (Promega, catalog no. A6120). Zika virus was detected using primer pair: forward 5′-CCGCTGCCCAACACAAG-3′ and reverse 5′-CCACTAACGTTCTTTTGCAGACAT-3′. The probe sequence is 5′-AGCCTACCTTGACAAGCAGTCAGACACTCAA-3′ 6-FAM/ZEN/IBFQ (Foy et al., 2011, Lanciotti et al., 2008). The cycling conditions involved activation at 45°C for 15 min and 95°C for 2 min, followed by 40 amplification cycles of 95°C for 15 s, and 60°C for 1 min.

Viral RNA was quantified by comparing each sample’s threshold cycle (*CT*) value with a ZIKV RNA standard curve.

### Western blotting

For protein extraction, cells were lysed on ice in cell lysis buffer (1% Triton X-100, 0.05% SDS, 100 mM Na2 PO4, and 150 mM NaCl) supplemented with protease inhibitor mixture (Roche) and PMSF for 15 min followed by centrifugation at 16,000 *g* for 15 min at 4 °C to remove cell debris. The protein concentration was determined by bicinchoninic acid (BCA) assay (Pierce, catalog no. 23223, Rockford, IL). 30 μg of each protein lysate were electrophoresed on a 12% SDS-polyacrylamide gel. The proteins were then transferred onto polyvinylidene difluoride membranes (EMD Millipore). The membrane was blocked with PBS-0.05% Tween 20 (PBS-Tween) containing 5% nonfat milk (Fisher Scientific, Pittsburgh, PA), and the membranes were washed three times and incubated with primary antibody in 1% milk PBS-Tween at 4 °C overnight. The membranes were then washed with PBS-Tween three times followed by a secondary antibody in 1% milk PBS-Tween for 2 h at RT. Immunoreactivity was detected using enhanced chemiluminescence (NEN Life Sciences, Waltham, MA) and imaged by Kodak 20000MM Image Station. Antibodies were diluted as following: 1:1000 anti-ISG20 (Proteintech, catalog no. 22097-1-AP); 1:10,000 anti-β-actin (Sigma, catalog no. A2066); 1:10,000 peroxidase-conjugated anti-rabbit IgG (Cell Signaling Technology, catalog no. 7074).

### IFNβ secretion analysis by ELLA

The IFNβ secretion of supernatants from the trophoblast cells with or without ZIKV infection was determined using the Simple Plex immunoassay system (ELLA, Protein Simple, San Jose, CA) as previously described (Aldo et al., 2016a). Briefly, 50 μl of sample was added to sample inlet ports on a cartridge, then the sample was split into multiple parallel channels from the sample inlet port. Each channel was specific for one particular analyte and subjected to a typical sandwich immunoassay protocol. The entire immunoassay procedure was automated, and the analyzed results were obtained using the manufacture-encoded calibration curves.

### Statistical analysis

Statistical analyses were performed using Prism software, version 8 (GraphPad, San Diego, CA). All data are presented as means ± SEM. Differences between two groups were analyzed using unpaired Student’s t-test and differences among multiple groups were analyzed by one-way ANOVA. Depending on the distribution of the continuous variables, nonparametric test was used if the data was not normally distributed. A *p*-value < 0.05 was considered statistically significant.

### Study approval

This study was carried out in accordance with the recommendations in the Guide for the Care and Use of Laboratory Animals of the National Institutes of Health. The protocols were approved by the Institutional Animal Care and Use Committee at the Wayne State University School of Medicine (Assurance number A3310-01). For the human primary trophoblast cell culture, a signed written consent form was obtained from the patients. The use of placental tissues, specimens and consent forms was approved by the Yale University Human Investigation Committee (#2000021607).

## Author Contributions

Conceptualization, G.M., P.A., and J.D.; Methodology, C.M.R., J.D., P.S., and Y.Y.; Investigation, J.D., H.L., X.Q., and J.J., Resources, B.L., D.B., A.L. and P.S.; Writing-Original Draft, J.D.; Writing-Review & Editing, C.M.R., P.S., A.M. and G.M.; Supervision, G.M., D.B., and A.L.; Funding Acquisition, G.M.

## Acknowledgement

This work was supported in part by NIH grant NIAID 1R01AI145829-01.

## Declaration of Interests

The authors have declared that no conflict of interest exists.

---

**Supplementary Figure 1.**
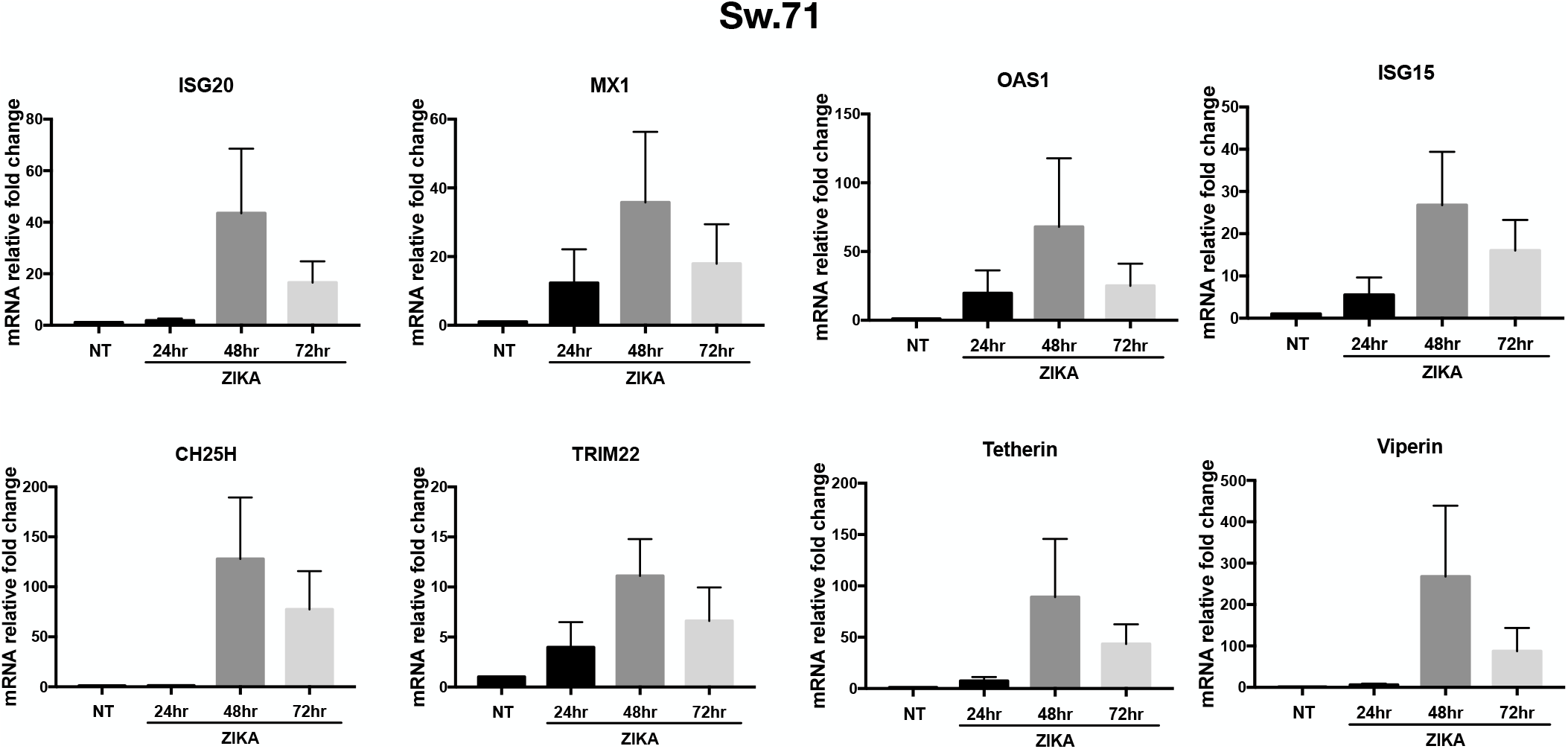
ZIKV infection results in a time-dependent interferon-stimulated gene (ISG) response in trophoblast cell line Sw.71. Human first-trimester trophoblast cell line Sw.71 cells were infected with ZIKV (MOI=2) for 1 h and refreshed with regular media over time. RNA was collected for gene expression by qRT-PCR. Data represent as mean ± SEM from three independent experiments. NT, no treatment group. Note that ISGs (*ISG20, MX1, OAS1, ISG15, CH25H, TRIM22, Tetherin, Viperin*) mRNA expressions were all induced in trophoblast cell line Sw.71.

**Supplementary Figure 2.**
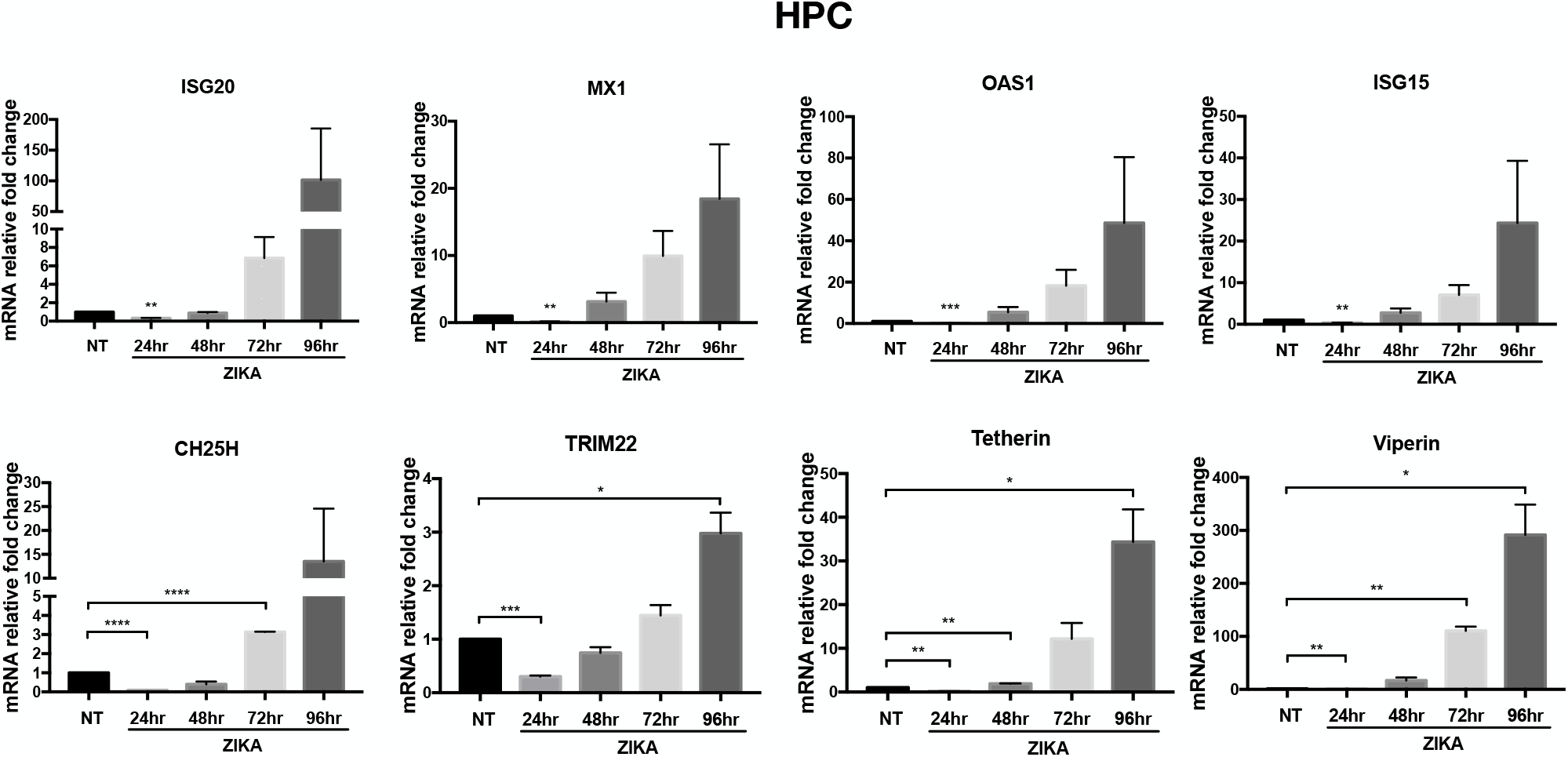
ZIKV infection results in a time-dependent interferon-stimulated gene (ISG) response in human primary culture trophoblast cells. Human primary culture trophoblast cells were infected with ZIKV (MOI=2) for 1 h and refreshed with regular media over time. RNA was collected for gene expression by qRT-PCR. Data represent as mean ± SEM from three independent experiments. \**p* < 0.05, \*\**p* < 0.01, \*\*\**p* < 0.001, \*\*\*\**p* < 0.0001. NT, no treatment group; HPC, human primary culture. ZIKV infection induced ISGs (*ISG20, MX1, OAS1, ISG15, CH25H, TRIM22, Tetherin, Viperin*) mRNA expression in human primary trophoblast cells. Note that ISGs expression were firstly inhibited in the first 24 h.p.i. and then increased afterwards.

**Supplementary Figure 3.**
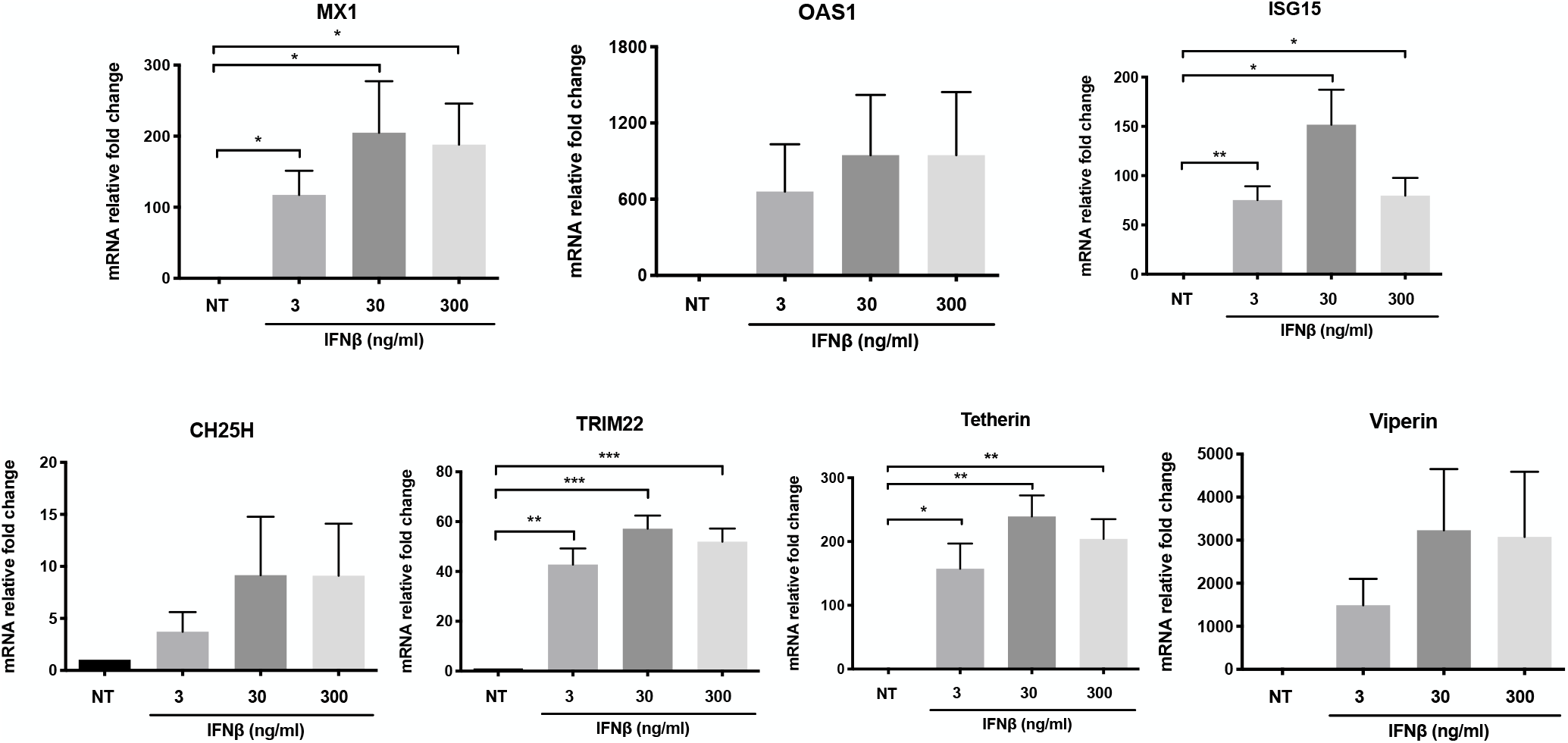
ZIKV-induced ISGs are the downstream of IFNβ signaling in trophoblast cells. Sw.71 cells were treated with different doses of IFNβ (3, 30, 300 ng/ml) for 8 h, and RNA was collected for determining gene expressions by qRT-PCR. Data represent as mean ± SEM from three independent experiments. \**p* < 0.05, \*\**p* < 0.01 and \*\*\**p* < 0.001. Note that even at the lowest dose (3 ng/ml), IFNβ was able to induce a robust increase of ISGs mRNA expression.

**Supplementary Figure 4.**
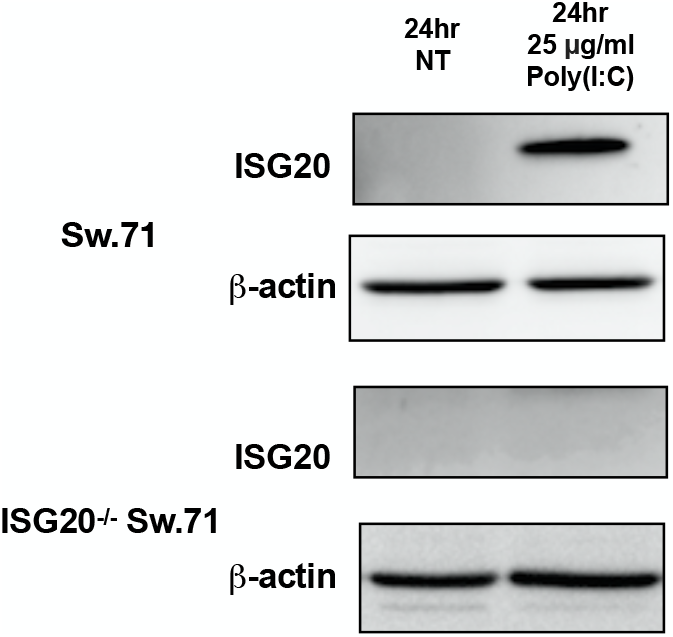
Verification of ISG20^−/−^ Sw.71 cells established by CRISPR/Cas9 system. Sw.71 and ISG20^−/−^ Sw.71 cells were treated with 25 μg/ml Poly(I:C) for 24 h, and proteins were collected for western blot analysis. β-actin served as a loading control. Note that no ISG20 protein expression was present in ISG20^−/−^ Sw.71 cells after Poly(I:C) treatment.

**Supplementary Figure 5.**
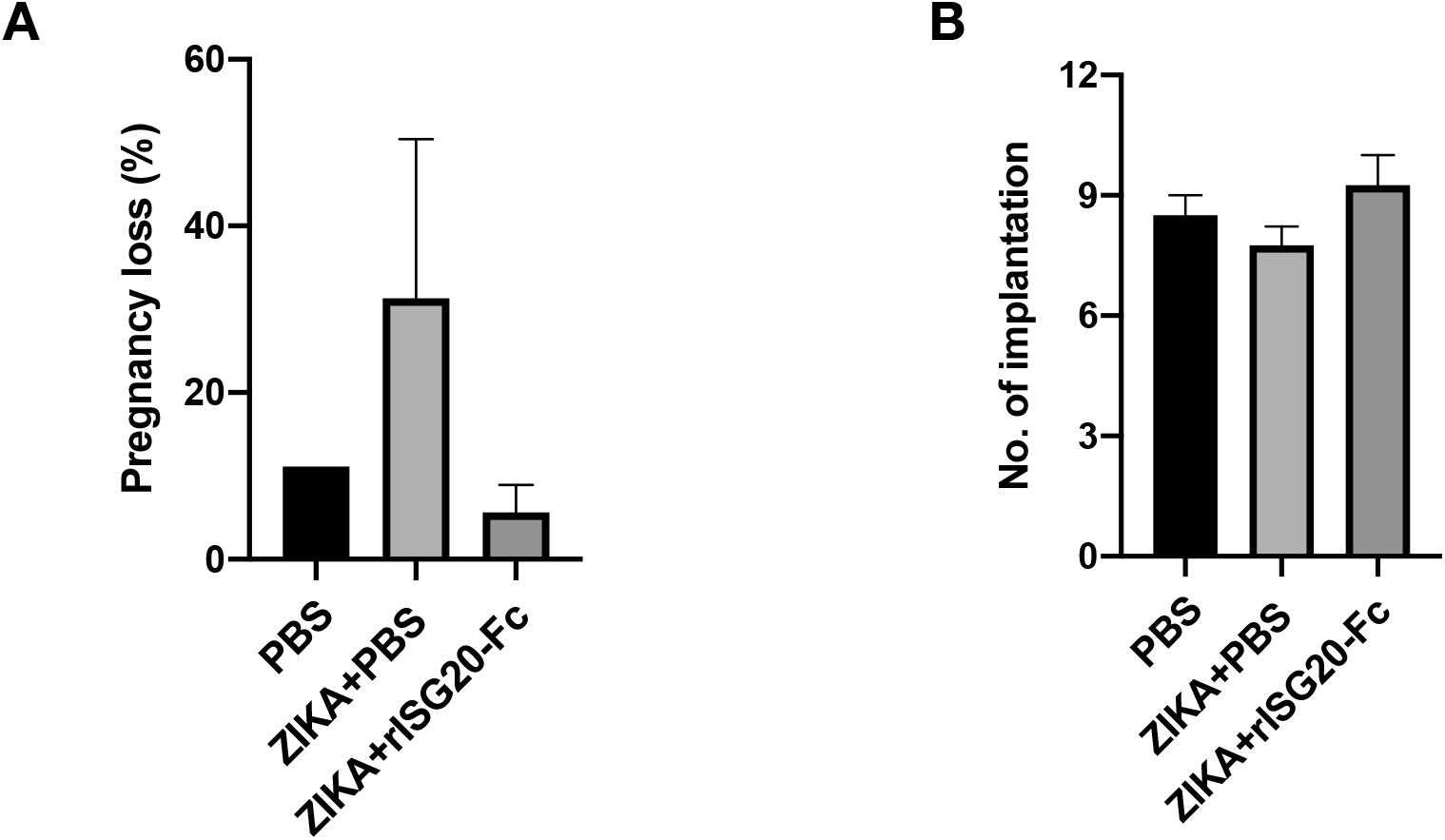
rISG20-Fc treatment reduces pregnancy loss in ZIKV-infected IFNAR1^−/−^ pregnant mice. **(A)** Effect of rISG20-Fc treatment on pregnancy loss. rISG20-Fc treatment decreased the pregnancy loss caused by ZIKA infection. The pregnancy loss rate (number of dead fetus + resorptions/ total number of implantations×100) was calculated. **(B)** Effect of rISG20-Fc treatment on implantation number. No differences were found in terms of the implantation number between groups. N=2 for control PBS group, and n=3-4 for treatment groups.

**Supplementary Table 1.**
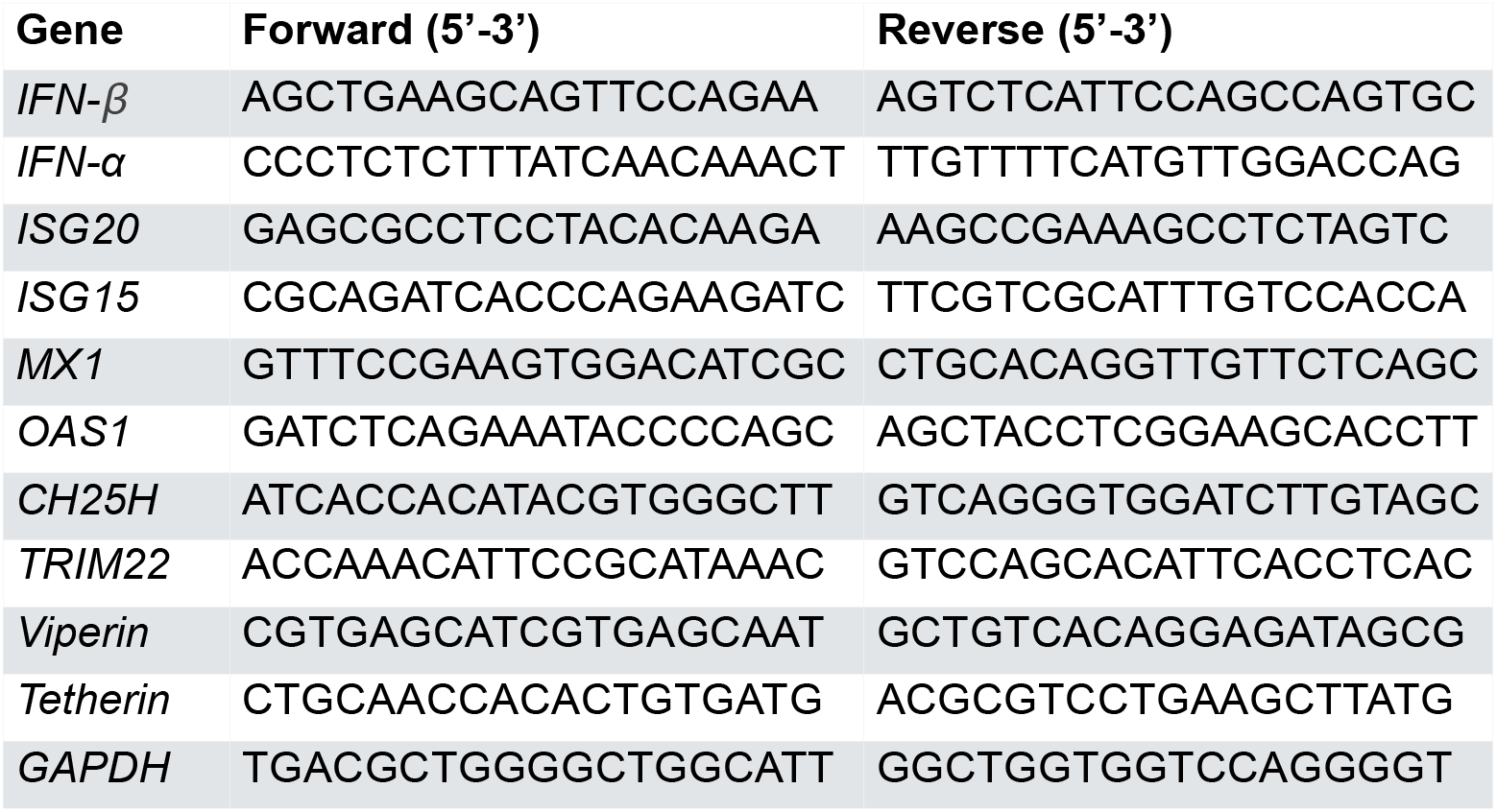
Primers used for qRT-PCR analyses.

